# RNA Binding Proteins KhpA and KhpB Interact with Small Regulatory RNAs and Affect Global Gene Expression in *Deinococcus radiodurans*

**DOI:** 10.1101/2025.09.18.676964

**Authors:** Runhua Han, Blanca I. Quinones-Diaz, Antonio Cordova, Brandon Niese, Phillip Sweet, Jaden Fang, Sean Engels, Angela Chen, Vernita D. Gordon, Lydia M. Contreras

## Abstract

Small regulatory RNAs (sRNAs) in bacteria often associate with RNA-binding proteins to gain intracellular stability and/or to enable regulatory efficiency. While much of the current knowledge about those sRNA binding proteins is derived from studies in Gram-negative organisms, the characterization of such proteins in Gram-positive species is still lagging behind. Here, we identified and characterized two sRNA binding proteins (KhpA and KhpB) in *Deinococcus radiodurans*, a Gram-positive bacterium that exhibits extreme resistance to radiation and other oxidative stressors. We demonstrate that KhpA and KhpB interact with key sRNAs in *D. radiodurans* and influence their stabilities. Although KhpA and KhpB interact with each other, they do not bind the sRNAs as a complex. Insightfully, KhpA and KhpB facilitate the interactions of the representative sRNAs *PprS* and *Dsr9* in *D. radiodurans* with their respective mRNA targets, *pprM* and *DR_1968*. Through RNA-seq analysis, we further revealed that KhpA and KhpB have both overlapping and specific roles in a global gene regulation in *D. radiodurans*. Overall, this study expands our knowledge of posttranscriptional regulation in *D. radiodurans* and supports the growing consensus that KhpA and KhpB homologs constitute a new family of sRNA binding proteins in Gram-positive bacteria.

**IMPORTANCE:** The bacterium *Deinococcus radiodurans* is the most radiation-resistant organism identified to date. Understanding the mechanisms of resistance of *D. radiodurans* is essential for leveraging this bacterium in biomedical and biomanufacturing applications. It was previously revealed that small regulatory RNAs (sRNAs) play crucial roles in the gene regulation of *D. radiodurans*. However, how these sRNAs are influenced by RNA binding proteins is poorly understood. Here we identified two conserved RNA binding proteins, KhpA and KhpB, as sRNA binding partners in *D. radiodurans*. These proteins affect the sRNA stability, sRNA-target interaction, and global gene regulation. Characterization of KhpA and KhpB will help us advance the understanding of how post-transcriptional network regulates the physiology and radioresistance of *D. radiodurans*.

## INTRODUCTION

Small regulatory RNAs (sRNAs) act as regulators of gene expression in nearly all bacteria (1). In-depth studies on sRNAs over the past 20 years have revealed the existence of large post-transcriptional networks that rival the complexity of primary gene expression control and have an effect on every aspect of cellular function (2). Most sRNAs influence the regulation of mRNA translation and/or stability through antisense binding to complementary regions in mRNA targets (2–4). Importantly, interaction with certain RNA binding proteins (RBPs) can assist the base-pairing between sRNAs and their target mRNAs before, during, or after the sRNA-target interactions (3, 4). RBPs can facilitate recognition between the two interacting RNAs, promote RNA folding by loosening their structures, or induce specific degradation/hydrolysis of the sRNA/target RNA (3).

Three proteins–Hfq, CsrA, and ProQ–have been characterized as sRNA-associated RBP in model bacteria. Among these proteins, Hfq is the most well-studied and is present in approximately 50% of bacterial species (3). Although not required for the structural rearrangement of all sRNA targets (5), Hfq acts as a chaperone by stabilizing sRNAs and/or as a matchmaker that promotes interactions between sRNA and mRNA pairs by inducing changes in the secondary structures of sRNAs (6). For instance, Hfq binds to the *dgcM* mRNA in *Escherichia coli*, unfolding inhibitory structures and making *dgcM* more accessible as a target to its cognate sRNAs, OmrA and OmrB (7). A second widespread class of bacterial sRNAs interacts with the protein CsrA. CsrA has been reported to bind various sRNAs across different bacteria, such as CsrB/C, FnrS, SgrS and McaS in *E. coli* and FliW in *Bacillus subtilis*). These sRNAs sequester CsrA, inhibiting its RNA-binding activity and regulating hundreds of mRNAs targets of the protein (8, 9). Interestingly, CsrA has also been recently implicated in regulating sRNA-mRNA interactions (10). In recent years, ProQ was identified as an additional class of bacterial sRNA binding proteins. ProQ promotes base-pairing between sRNAs and their target mRNAs by recognizing the secondary structures on the 3’ end of the mRNAs (11, 12). *In vivo* studies, such as UV Crosslinking and RNA Sequencing (CLIP-seq) and RNA Interaction by Ligation and Sequencing (RIL-seq), have mapped hundreds of ProQ binding sites on mRNAs and sRNAs in *E. coli* and *Salmonella Typhimurium* (12). Intriguingly, a significant portion of the ProQ-bound RNA pairs was found to be also associated with Hfq, suggesting the overlapping, complementary, or competing roles of the two proteins (13).

Importantly, while many prokaryotes encode for functional sRNAs, some lack homologs of Hfq, ProQ, or CsrA. This strongly suggests the existence of other families of RNA-binding proteins that may affect the sRNA functions and perhaps contribute to additional cellular functions. Notably, two KH-domain containing proteins, KhpA and KhpB, have been identified as global RNA binding proteins in a number of gram-positive bacteria, including *Streptococcus pneumoniae*, *Clostridioides difficile*, *Lactobacillus plantarum, Enterococcus faecalis/faecium* and *Fusobacterium nucleatum* (14–17). RNA co-immunoprecipitation sequencing (RIP-seq) have revealed that KhpA and/or KhpB are able to bind sRNAs, tRNAs, and mRNAs, though their binding preferences and RNA distributions vary among species (14, 16–18). Furthermore, KhpA and KhpB were found to physically interact with each other in *S. pneumoniae* and *E. faecalis/faecium* (14, 16). Many RNAs were co-purified with one or both proteins, suggesting cooperative roles for KhpA and KhpB in binding and regulation of these RNAs. However, although deletion of *khpA* or *khpB* resulted in altered mRNA expression and changed sRNA stability in *C. difficile* and *F. nucleatum* (17, 18), the broader impact of KhpA and KhpB on global gene expression in other bacterial species is still largely unknown. Nevertheless, KhpA and/or KhpB have been implicated in cell division and growth in all the organisms in which these proteins have been studied thus far (14, 15, 17, 18). The sRNA binding ability of KhpA and KhpB, coupled with their conservation across many bacteria (19) strongly suggests that they might fulfill the role of sRNA chaperones in these species. However, the mechanism by which KhpA and KhpB modulate sRNA-target interaction and the conservation of their functions across gram-positive bacteria remain poorly understood. In addition, the biological importance of the KhpA-KhpB interaction in RNA biology remains an open question.

*Deinococcus radiodurans* is renowned for its exceptional ability to withstand high levels of oxidative stresses such as ionizing radiation, UV light, and desiccation (20). The study of *D. radiodurans* as a model organism for oxidative stress resistance holds significant importance for medicine and public health. The exceptional phenotype of *D. radiodurans* can be attributed to a number of factors, including: i) a highly effective and redundant DNA repair system; (ii) the presence of numerous antioxidant metabolites that enhance proteome protection; and (iii) a highly condensed nucleoid structure (21–23). Importantly, the posttranscriptional regulation mediated by sRNAs has been shown to play important roles in *D. radiodurans*’s response to oxidative stress (24–28). For example, the well-characterized sRNA in *D. radiodurans*, *PprS*, promotes survival of the bacterium under ionizing radiation by stabilizing *DR_0907*/*pprM*, a modulator of pleiotropic proteins involved in DNA repair (29). It is worth noting that KhpA and KhpB in *D. radiodurans* (*DR_2009* and *DR_0246*, respectively) have been linked with genome condensation phenotypes associated with oxidative stress responses (30). The potential functional importance of KhpA and KhpB to *D. radiodurans* is further evidenced by their conservation across other *Deinococcus* species (19), which lack homologs of Hfq, ProQ, and CsrA. Despite this, the specific physiological roles of KhpA and KhpB, as well as their involvement in sRNA mediated regulation in *D. radiodurans* is unclear.

In this study, we experimentally confirm that KhpA and KhpB function as sRNA-binding proteins in *D. radiodurans*. We demonstrate that KhpA and KhpB resemble the activities of canonical sRNA binding proteins (e.g., Hfq and ProQ) by binding key sRNAs, affecting their stability and interaction with mRNA targets. The deletion of *khpA* and *khpB* also affects expression of genes involved in various pathways, many of which were known to be regulated by KhpA/B interacting sRNAs. Overall, these findings support recent observations that KhpA and KhpB homologs represent a new class of bacterial RBPs with sRNA binding and regulation functions.

## RESULTS

### Identification of sRNA binding proteins in *Deinococcus radiodurans*

To identify the proteins that potentially interact with sRNAs in *D. radiodurans*, we first carried out an MS2-sRNA affinity purification to capture the protein partners of *PprS*, a well-characterized sRNA in *D. radiodurans* (29). In this experiment, the MS2 aptamer was fused to the 5’ end of *PprS*. The MS2-*PprS* fusion was expressed from the pRADgro plasmid in the *D. radiodurans* wild-type strain (WT) and purified during the mid-exponential growth phase. The co-purified proteins (**Figure S1A**) were subsequently identified by LC-MS/MS followed by bioinformatic analysis (**Figure 1A**). By comparing normalized read counts of the MS2-*PprS* and the control sample (untagged PprS expressing from the pRADgro plasmid), 16 proteins were significantly enriched (fold change ≥ 1.5, adjust *p* ≤ 0.05) in the MS2-*PprS* pull-down compared to the control (**Figure 1B, Table S1**). These included the DNA repair protein PprA, an RNA 2’,3’-cyclic phosphodiesterase, the 30S ribosomal protein, and several metabolic and hypothetical proteins. It is worth noting that DR_2009/KhpA emerged as one of the top hits and was the only identified protein with a known RNA binding domain.

**Figure 1.**
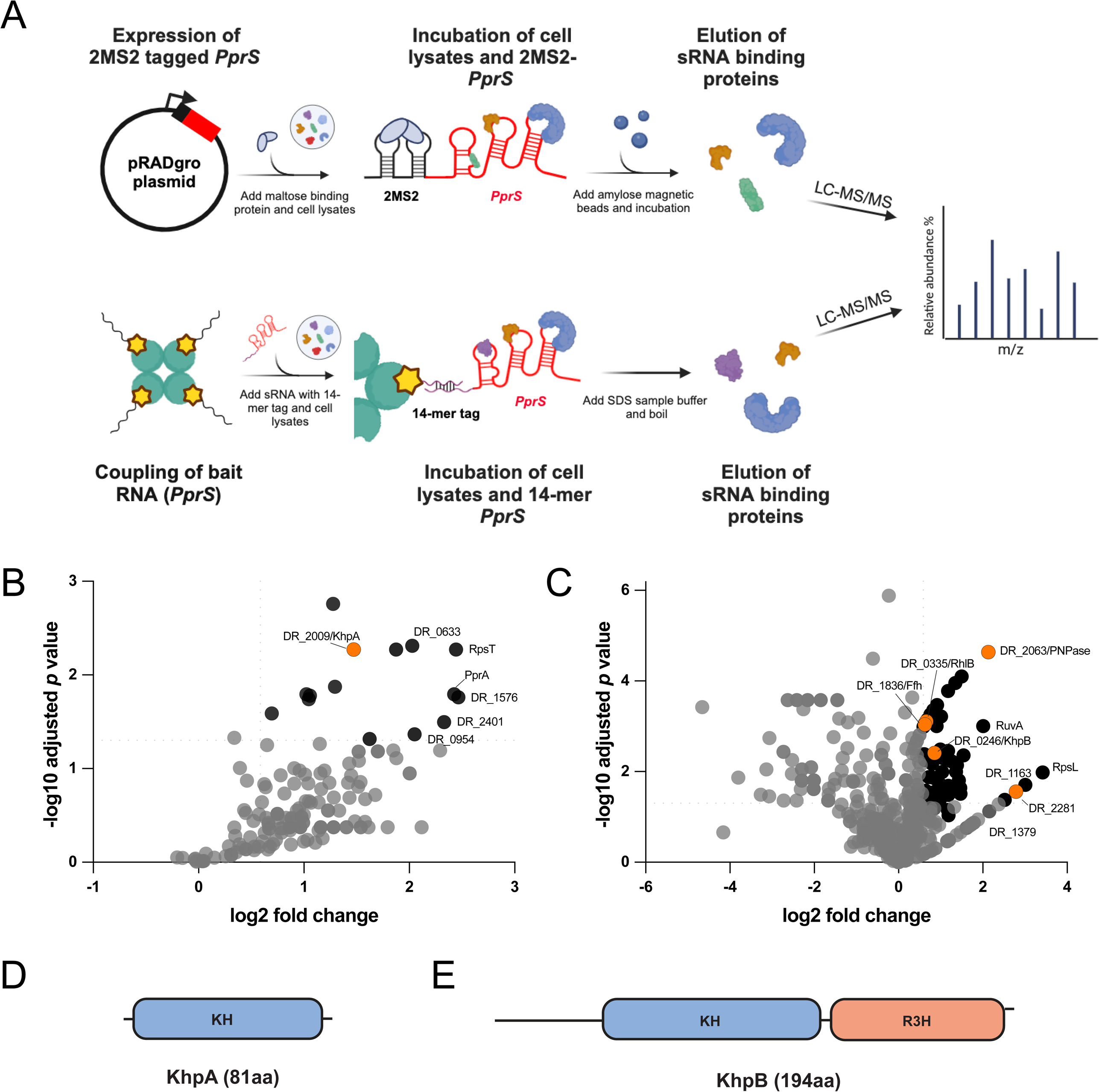

Given earlier reports indicating that MS2-sRNA affinity purification can be less efficient in some gram-positive bacteria (i.e., *F. nucleatum and S. pneumoniae*) (17, 31), we considered the possibility that this method could fail in capturing certain sRNA interacting RBPs in *D. radiodurans*. To address this limitation, we employed a second protocol originally developed for identifying eukaryotic miRNA interacting RBPs (32). In this assay, the sRNA (*PprS*) and a control randomized RNA of the same length was transcribed *in vitro* with a 5’-located 14-nt long tag and incubated with lysates extracted from *D. radiodurans* cells in the mid-exponential phase. The resulting co-purified proteins (**Figure S1B**) were subjected to LC-MS/MS (**Figure 1A**). Using this alternative approach (hereinafter referred to as 14-mer pull-down), we successfully enriched a total of 66 proteins (fold change ≥ 1.5, adjust *p* ≤ 0.05) in the 14-mer *PprS* sample compared to the control (**Figure 1C, Table S1**). Among the enriched proteins, 27 were predicted to have potential nucleic acid binding functions; these included three ribosomal proteins, two translation initiation factors, two TetR transcriptional regulators, five tRNA modification/processing proteins, DNA polymerase I, and five DNA repair proteins (e.g., RuvB, SbcD, and RecR). Notably, DR_2281, which contains a double stranded RNA binding motif, was one of the most enriched RBPs, although its specific function remains unknown. Five other proteins (PNPase, RhlB, Ffh, DR_2281, KhpA and KhpB) were identified as RBPs with known RNA binding domains. PNPase, RhlB, and Ffh are known to play crucial roles in oxidative stress response in *D. radiodurans* by interacting with either oxidized RNA or Signal Recognition Particle RNA (25, 33). PNPase also plays an indispensable role in paradoxically stabilizing sRNAs bound to other RBPs (e.g., Hfq) and promoting sRNA-mediated in gene regulation in *E. coli* (indicating a similar role in *D. radiodurans*) (34). In contrast, although KhpA and KhpB have been demonstrated to interact with sRNAs in other bacterial species, their specific contributions to sRNA regulation have not been studied in *D. radiodurans*. As such, we focused on KhpA and KhpB and further characterized their roles in *D. radiodurans*.

### KhpA and KhpB are highly conserved RNA-binding proteins within the genus Deinococcus

In *D. radiodurans*, KhpA is a small protein consisting of 81 amino acids and contains a single-KH RNA binding domain (**Figure 1D**), which is a well-characterized structure that interacts with RNA backbone residues through a conserved GXXG motif (19). KhpB, on the other hand, has two RNA binding domains, KH and R3H, at the C terminus (**Figure 1E**). The RNA-binding domains of KhpA and KhpB in *D. radiodurans* share significant similarity with their homologs in other bacteria, such as *S. pneumoniae*, *C. difficle,* and *F. nucleatum* (**Figure S2A-S2B**). However, unlike its counterparts in these species, KhpB in *D. radiodurans* lacks the Jag-like domain at the N terminus (14, 16–18). Phylogenetic conservation analysis revealed that KhpA and KhpB homologs are highly conserved across publicly available *Deinoccoccus* genomes, exhibiting 92-100% sequence identity for KhpA and 95-100% for KhpB compared to their *D. radiodurans* counterparts (**Figure S2C-S2D**). Interestingly, the gene synteny of KhpA and KhpB in *D. radiodurans* is distinct from their homologs in other bacteria. While *khpA* and *khpB* in *C. difficile*, *S. pneumoniae* and *F. nucleatum* are adjacent to upstream genes encoding for small ribosomal subunit protein bS16 (RpsP) and membrane protein insertase YidC/Oxa1, KhpA and KhpB in *D. radiodurans* are syntenic with upstream genes encoding aspartate-semialdehyde dehydrogenase (Asd) and acyl-carrier-protein synthase (AcpS), respectively (**Figure S2E-S2F**). This unique gene arrangement raises the possibility that these two proteins in the bacterium may have expanded or distinct functionalities in *D. radiodurans*. Despite these differences, the high conservation of their RNA binding domains strongly suggests that KhpA and KhpB in *D. radiodurans* likely function similarly to their homologs in other bacteria, particular with regard to sRNA binding.

### KhpA and KhpB singly interact with important sRNAs in *D. radiodurans*

To confirm the association of KhpA and KhpB with the *PprS* sRNA observed in the RNA affinity pulldown experiments (**Figure 1B-1C**), and to further investigate their sRNA binding capabilities, we performed electrophoretic mobility shift assays (EMSA) by incubating purified KhpA or KhpB protein (**Figure S3A**) with radiolabeled *PprS*. As shown in **Figure 2**, both KhpA and KhpB were able to form stable complexes with *PprS*. By calculating the dissociation constant (K_d_), we found that KhpA exhibits a higher binding affinity (972.7 ± 83.5 nM) for *PprS* compared to KhpB (5605.4 ± 303.1 nM) (**Table 1**). To examine whether this interaction was specific to *PprS*, we tested three additional sRNAs (*Dsr9*, *Dsr11*, and *Dsr20*) that were confirmed to play important roles in stress response of *D. radiodurans* (35). All three sRNAs exhibited concentration-dependent shifts when incubated with KhpA and KhpB (**Figure 2**), suggesting that the interaction was not unique to *PprS*. Consistent with the *PprS* data, the sRNA binding affinities of KhpA are generally higher than those of KhpB (**Table 1**). Among the tested sRNAs, *Dsr11* displayed the strongest binding affinity for both KhpA and KhpB (**Table 1**), although the binding strengths for all sRNAs were in the same order of magnitude. It is worth noting that our observed sRNA binding affinity of KhpA and KhpB is similar to those reported in *F. nucleatum* (both in the micromolar range) (17), but significantly lower than the low nanomolar affinities observed for canonical sRNA binding proteins like Hfq, or CsrA (8, 36, 37). This indicates that KhpA and KhpB are relatively weaker sRNA binders. Another intriguing observation is that KhpA forms two complexes with all four tested sRNAs, as reflected by the supershifts (indicated by the green arrows in **Figure 2A-2D**), whereas KhpB forms a single complex (indicated by the orange arrows in **Figure 2E-2H**). This observation agrees with a previous study showing that KhpA can form a homodimer itself (38), suggesting that each sRNA can bind two KhpA molecules. Importantly, no complexes were detected when a randomized RNA (with a random sequence) was incubated with KhpA or KhpB (**Figure S3B-S3C**), implying that the observed interactions between these proteins and sRNAs were not artificial.

**Figure 2.**
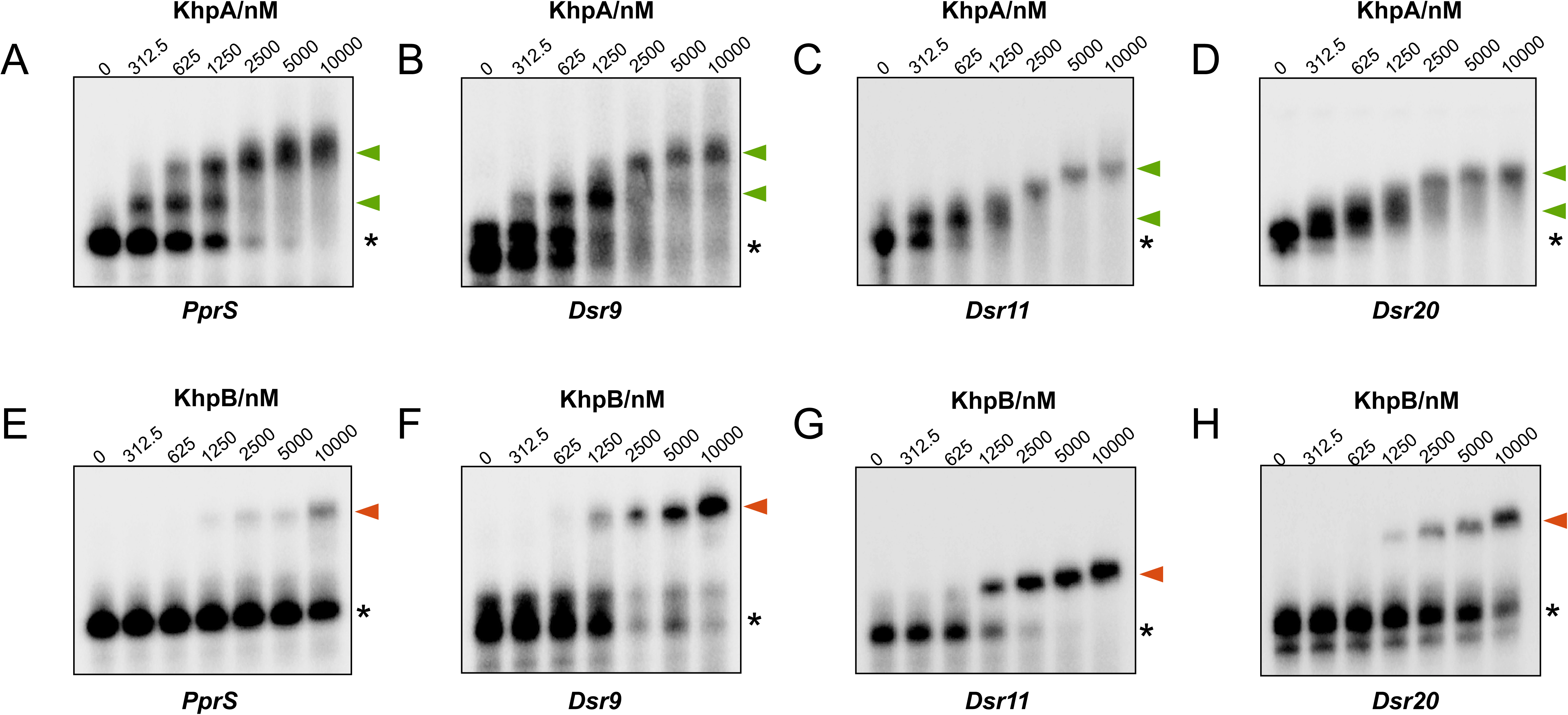

**Table 1.** Binding affinity of KhpA and KhpB on selected sRNAs in *D. radiodurans*. The dissociation constants (K_d_) were determined in duplicate by EMSA (Figure 2).

| sRNA | Stress response | Representative target | $K_d$/KhpA, nM | $K_d$/KhpB, nM |
| --- | --- | --- | --- | --- |
| *PprS* | Ionizing radiation | *DR_0907/pprM* | 972.7±83.5 | 5605.4±303.1 |
| *Dsr9* | H<sub>2</sub>O<sub>2</sub> | *DR_1968* | 530.2±61.2 | 2913.1±129.0 |
| *Dsr11* | Heat | *DR_1615* | 282.1±19.2 | 885.1±63.1 |
| *Dsr20* | H<sub>2</sub>O<sub>2</sub> | *DR_1968* | 666.4±85.0 | 3312.8±346.1 |

### KhpA and KhpB affect the stability of *D. radiodurans* sRNAs

We next investigated if the binding of KhpA and KhpB influences the fate of these sRNAs. RNA decay experiments were then carried out to examine the stability changes of *PprS*, *Dsr9*, *Dsr11*, and *Dsr20* by Northern blotting or qPCR (**Figure 3A-3D, Figure S3D)**. These experiments revealed differences in the inherent stabilities of the tested sRNAs in WT *D. radiodurans*. *PprS* exhibits high stability, with little changes in abundance after the addition of rifampicin, while Dsr20 showed moderate stability. In contrast, Dsr9 and Dsr11 were relatively unstable, with their levels decreasing rapidly after the halt of transcription (**Figure 3A-3D**). Importantly, we also observed changed stabilities of these sRNAs in deletion strains lacking KhpA or KhpB. Upon the deletion of KhpA, the stability of *PprS* remained unchanged within 60 min after rifampicin treatment, but the half-lives of Dsr9 and Dsr20 were notably reduced in the *ΔkhpA* strain (**Figure 3E**). Dsr11 also showed a shorter half-life in the absence of KhpA; however, this change was not significant (**Figure 3E**). In the *ΔkhpB* strain, the half-lives of Dsr9 and Dsr11 in the Δ*khpB* strain were significantly prolonged compared to the WT strain. Conversely, the stability of Dsr20 was not significantly altered between the WT and Δ*khpB* strains (**Figure 3E**). Similar to Δ*khpA*, we did not observe any changed stability of *PprS* in Δ*khpB* within the tested timeframe. These data suggest that KhpA and KhpB both function as sRNA stability factors in *D. radiodurans*, but their effects are sRNA specific and functionally distinct: KhpA stabilizes certain sRNAs, whereas KhpB appears to have negative effects on sRNA stability.

**Figure 3.**
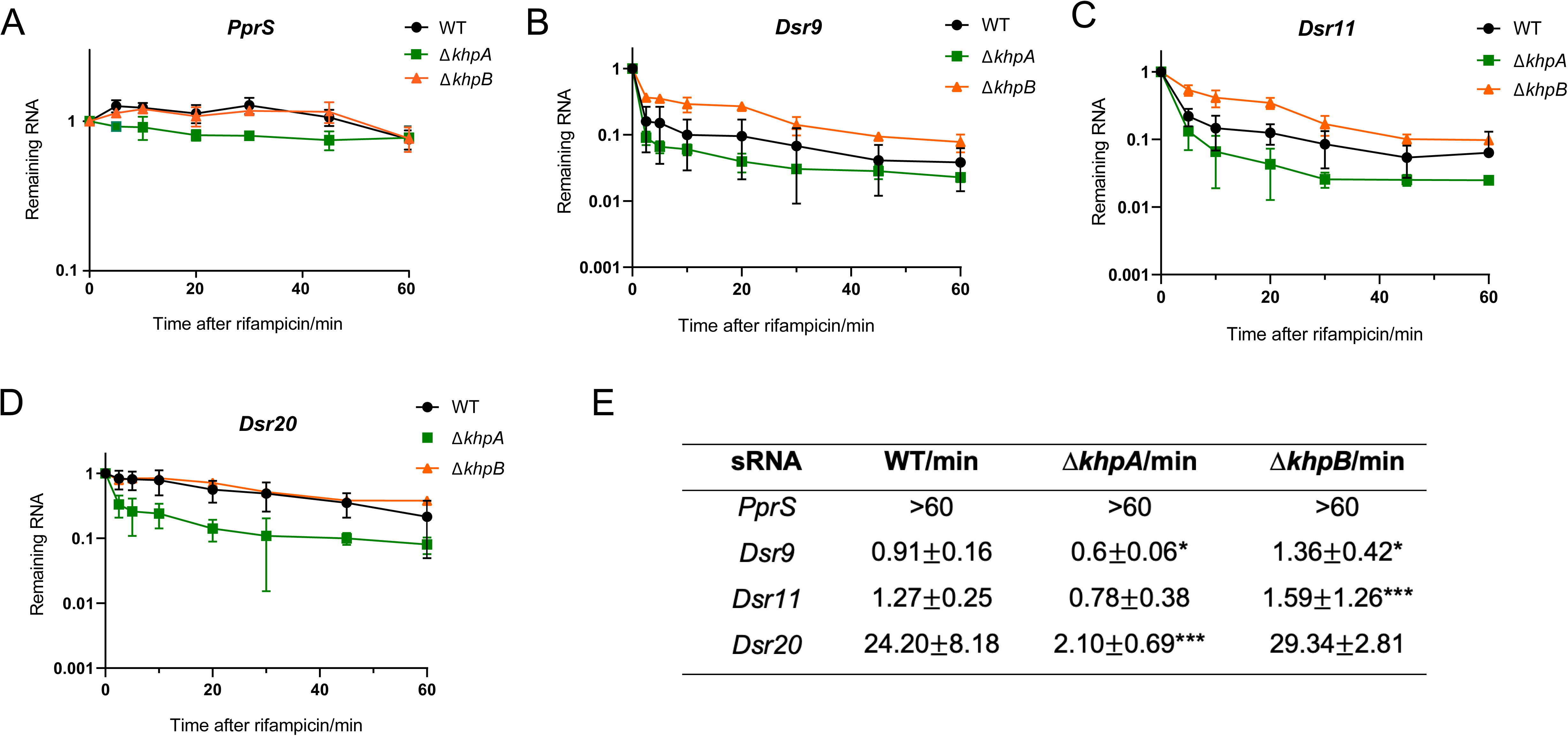

### KhpA and KhpB form complex but bind sRNA independently

It was previously shown that KhpA and KhpB form a complex in *S. pneumoniae* and *E. faecalis/faecium* (14, 16). To examine if KhpA and KhpB form a complex and if other proteins associated with KhpA or KhpB in *D. radiodurans*, we conducted protein-protein coimmunoprecipitation (coIP) experiments using 3xFLAG-tagged fusions as bait. KhpA was tagged at the C-terminus which expressed on the pRADgro plasmid (KhpA-3xFLAG), while KhpB was tagged at the C-terminus chromosomally (KhpB-3xFLAG). For controls, we used a Δ*khpA* strain expressing non-tagged KhpA (the KhpA complementation strain; KhpA Comp) for KhpA coIP, and the WT strain for KhpB coIP. The purified proteins (**Figure S1C**) were subjected to LC-MS/MS. Compared to KhpA Comp, only 8 proteins were enriched significantly with KhpA-3xFLAG (**Figure 4A**, **Table S2**). By contrast, 28 proteins were significantly enriched (fold change ≥ 1.5, adjust *p* ≤ 0.05) with KhpB-3xFLAG relative to the WT strain expressing untagged KhpB (**Figure 4B**, **Table S2**), implying that KhpB might be involved in larger protein networks relative to KhpA in *D. radiodurans*. These proteins include two tRNA ligases, the elongation factor G, a metal binding protein, four ribosome-associate proteins as well as several metabolic proteins. Another RBP, Rsr, was also co-purified with KhpB, indicating a functional interaction between the two proteins in *D. radiodurans*. Importantly, we were able to reciprocally co-purify KhpA and KhpB with each other, suggesting an active interaction between the two proteins in *D. radiodurans*. To further validate this, KhpB was tagged with 6xHis on the chromosome in the KhpA-3xFLAG background. The coIP experiments were repeated using KhpA-3xFLAG or KhpB-6xHis as a bait, followed by Western blotting analysis. As observed in **Figure 4C** and **4D**, KhpB-6xHis was successfully co-purified in the KhpA-3xFLAG-coIP, and vice versa. Notably, the co-purifications between KhpA and KhpB were not abolished by RNase treatment of the cell lysates (**Figure 4C-4D**), demonstrating that the interaction between KhpA with KhpB does not rely on their RNA substrates. An interaction between purified KhpA and KhpB proteins was also detected using size-exclusion chromatography (**Figure S3E**). Thus, a stable KhpA and KhpB interaction is present in *D. radiodurans* in an RNA-independent manner.

**Figure 4.**
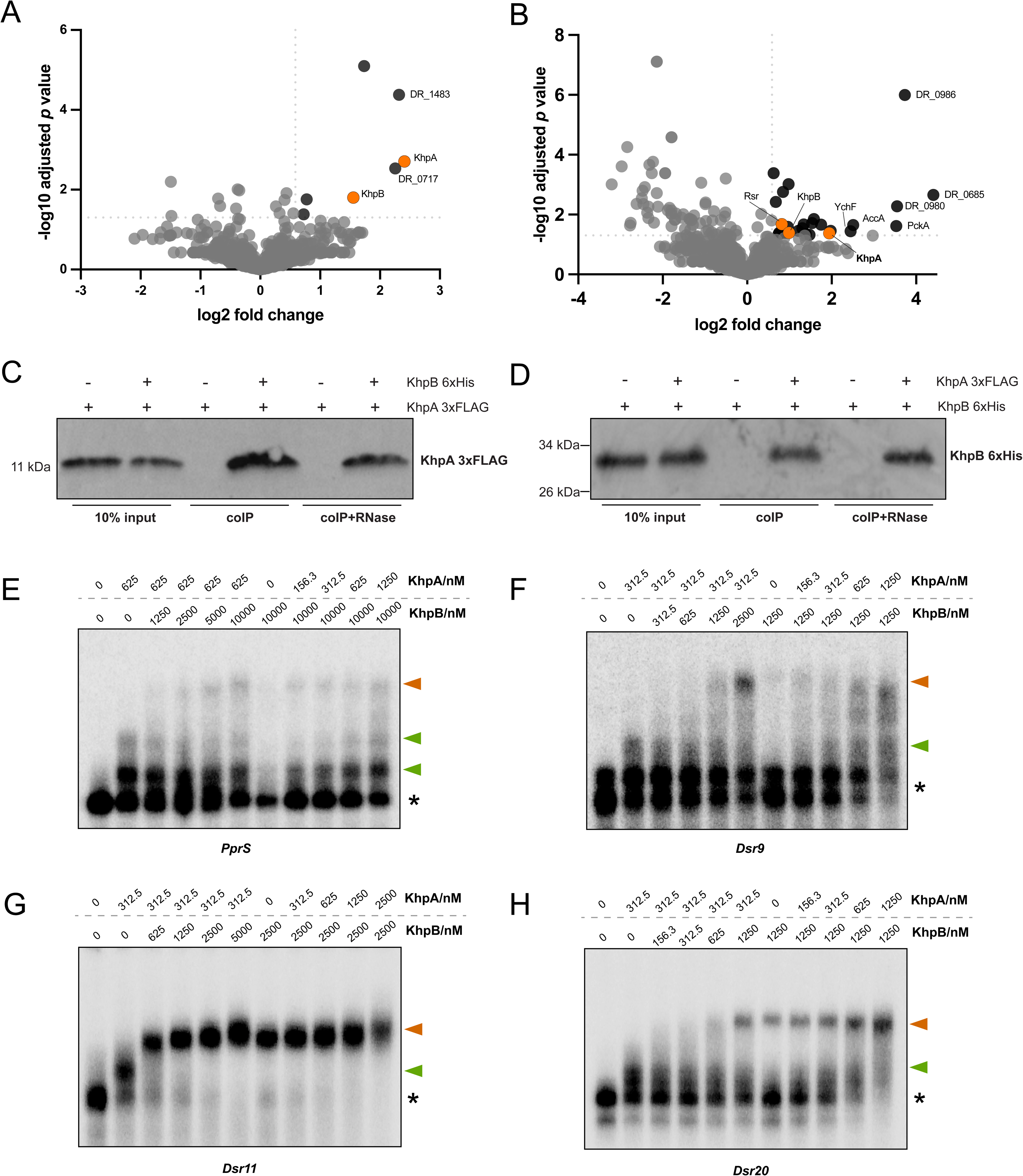

The co-occurrence of *khpA* and *khpB* genes in the genomes of many bacteria (19) and the interaction between the two proteins in *D. radiodurans* indicate that KhpA and KhpB may bind sRNAs as a heterocomplex, either alone or in complex with other proteins. To test this hypothesis, we incubated radiolabeled *PprS*, *Dsr9*, *Dsr11*, and *Dsr20* with both KhpA and KhpB proteins and examined the complex formation between these molecules via EMSA assay. As shown in **Figure 4E-4H**, distinct bands corresponding to KhpA- and KhpB-sRNA complexes were observed when the two proteins were added to the same reaction. However, no supershift indicative of a KhpA-KhpB-sRNA ternary complex with a larger size than the KhpB-sRNA complex was found in the gel, suggesting that KhpA and KhpB do not bind these sRNAs as a complex in *D. radiodurans*. Instead, they might bind to these sRNAs and exert their functions on these sRNAs individually.

### Effect of KhpA and KhpB on gene expression in *D. radiodurans*

To further investigate the global regulatory roles of KhpA and KhpB in *D. radiodurans*, we performed RNA-seq analysis on Δ*khpA* and Δ*khpB* strains growing at mid-exponential phase. The analysis reveals that expression levels of 286 and 303 transcripts were significantly affected by the deletion of *khpA* and *khpB*, respectively (log2 fold change ≥ 1 or ≤ −1, adjusted *p* value ≤ 0.05; **Table S3**). Importantly, the expression of downstream genes (*rimM* for *khpA* and *DR_0245* for *khpB*) was not significantly affected by the deletion of *khpA* and *khpB*, respectively. However, elevated expression of *rimM* or *DR_0245* was observed in the *khpA* or *khpB* complementation strain using qPCR (**Figure S4A**). These results indicate that, although the deletion of *khpA* or *khpB* did not cause any polar effect, they may play a role in regulating their downstream genes. To further validate the RNA-seq results, we confirmed the upregulation of one of the mostly affected genes in each respective deletion strain (*DR_B0060* and *DR_1571*, respectively) using qPCR. These changed expressions are consistent with the trends observed in the RNA-seq data (log2 fold change of ∼5.4 for *DR_B0060* in Δ*khpA* compared to WT, and log2 fold change of 5.4 for *DR_1571* in Δ*khpB*).These expression changes were partially restored to the WT levels in the corresponding complementation strains (**Figure S4B**). These data support the reliability of our RNA-seq data.

Among the differential expressed genes, a subset were affected by the deletion of both KhpA and KhpB (18.5% of KhpA-regulating genes and 17.5% of KhpB-regulating genes; **Figure 5A**; **Table 2**). These genes encode proteins with diverse functions, including the DNA damage response protein D (DR_0326), the HSP20 family heat shock protein (DR_1114), the Ro ribonucleoprotein Rsr (DR_1262), transcriptional regulators, and transporters. Interestingly, the changed expression of these transcripts showed similar trends upon KhpA or KhpB deletion, with only two exceptions **(Figure 5B**, **Table 2)**. GO Enrichment analysis (39) further revealed enrichment in pathways related to the (plasma) membrane, ATP binding and metal binding in both *khpA* and *khpB* knockout strains (**Figure 5C**). These data suggest the cooperation between the two proteins in the regulation of these genes and pathways. However, a larger number of transcripts were only affected by deletion of either *khpA* (233 transcripts) or *khp*B (250 transcripts). Interestingly, 84.5% (197/233) of the *khpA*-regulated genes were upregulated in the Δ*khpA* strain, whereas only 36.8% (92/250) of *khpB*-regulated genes showed this trend in the Δ*khpB* strain (**Figure 5A, Table S3**). This suggests that KhpA predominantly acts as a negative regulator of gene expression, while KhpB tends to serve a more positive regulatory role at the transcriptional level. The divergent regulatory effects were also reflected in cellular phenotypes: Δ*khpA* displayed larger nucleoids whereas Δ*khpB* cells tends to exhibit more condensed nucleoids (30). Further GO enrichment analysis highlighted distinct functional categories regulated by KhpA and KhpB. KhpA primarily influenced pathways related to ATP hydrolysis, DNA binding, siderophore-dependent iron import, and DNA integration (**Figure 5C**). In contrast, KhpB predominately affected signal transduction, proteolysis and phosphorylation pathways (**Figure 5C**). These findings suggest both overlapping and specific roles of KhpA and KhpB in global gene regulation in *D. radiodurans*.

**Figure 5.**
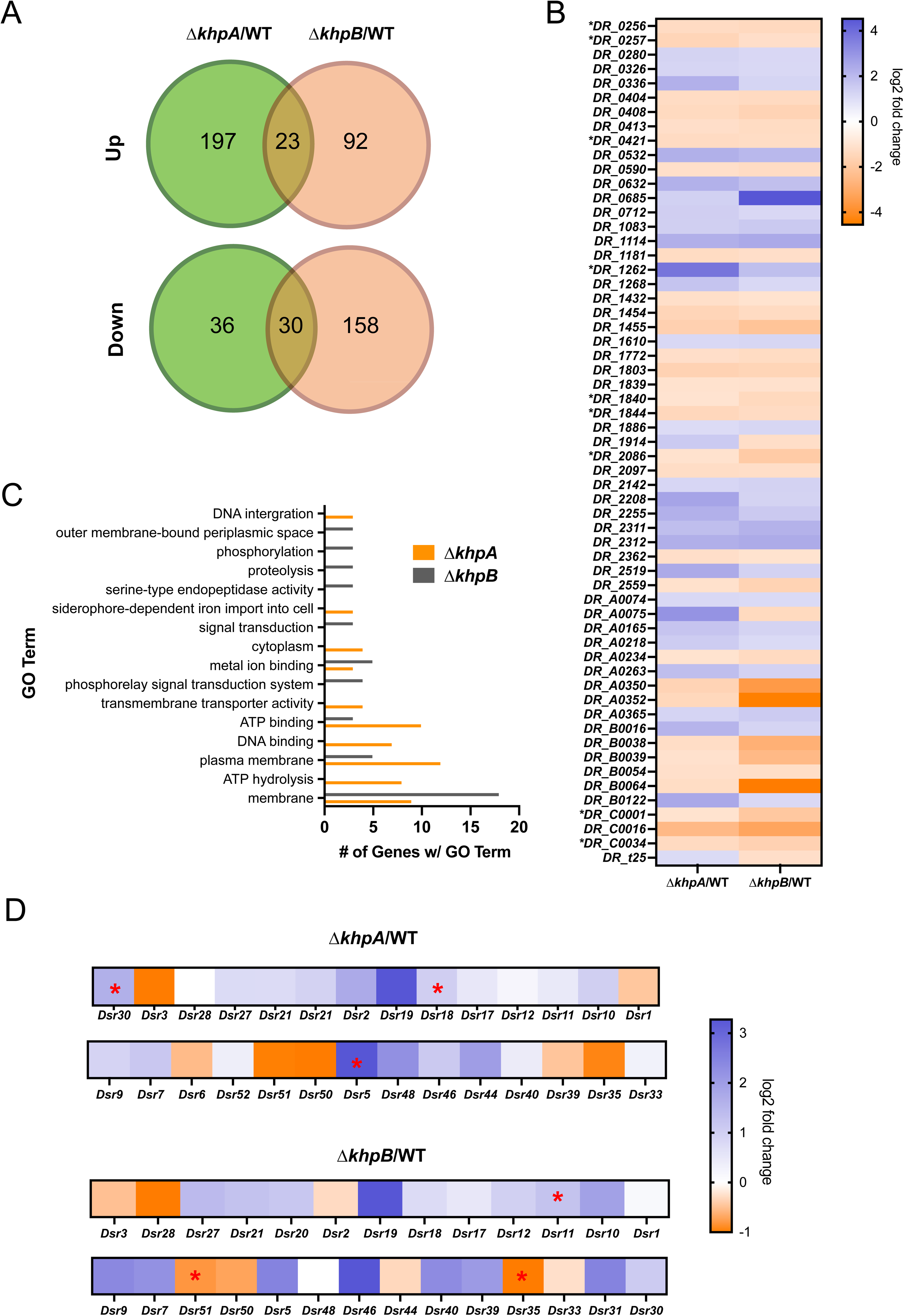

**Table 2.** Genes with differential expression in both Δ*khpA* and Δ*khpB* compared to WT. Known mRNAs bound to or affected by sRNAs in *D. radiodurans* are shown in bold, while genes displayed different trends in Δ*khpA* and Δ*khpB* are marked by asterisks.

| Gene Name | log2 foldchange<br>$\Delta khpA/WT$ | log2 foldchange<br>$\Delta khpB/WT$ | Function |
| --- | --- | --- | --- |
| **DR_0256** | -1.280149893 | -1.37293476 | Acetyltransferase component of pyruvate dehydrogenase complex |
| **DR_0257** | -1.503616218 | -1.161357467 | Pyruvate dehydrogenase E1 component |
| *DR_0280* | 1.242020899 | 1.116163409 | Branched-chain amino acid ABC transporter, periplasmic amino acid-binding protein |
| *DR_0326* | 1.183227034 | 1.098360397 | DNA damage response protein D |
| *DR_0336* | 2.203946537 | 1.197437024 | Fatty-acid--CoA ligase, putative |
| *DR_0404* | -1.256590513 | -1.338011894 | Uncharacterized protein |
| *DR_0408* | -1.348082929 | -1.580244868 | Response regulator |
| *DR_0413* | -1.164839607 | -1.27573939 | Uncharacterized protein |
| *DR_0421* | -1.298331982 | -1.232370566 | Uncharacterized protein |
| *DR_0532* | 2.218720816 | 1.998361783 | Uncharacterized protein |
| *DR_0590* | -1.131863219 | -1.257931436 | Uncharacterized protein |
| *DR_0632* | 2.198220238 | 1.791122135 | Uncharacterized protein |
| *DR_0685* | 1.319819139 | 4.52099016 | Uncharacterized protein |
| *DR_0712* | 1.368600987 | 1.117386561 | Uncharacterized protein |
| *DR_1083* | 1.351611243 | 1.541542643 | Uncharacterized protein |
| **DR_1114** | 2.226132908 | 2.394919309 | Heat shock protein, HSP20 family |
| *DR_1181* | -1.276790353 | -1.178262594 | Uncharacterized protein |
| **DR_1262** | 3.770025856 | 1.827854833 | Rsr |
| *DR_1268* | 1.671810603 | 1.099823061 | Uncharacterized protein |
| *DR_1432* | -1.156425714 | -1.034348834 | Uncharacterized protein |
| *DR_1454* | -1.514816183 | -1.307066838 | Uncharacterized protein |
| *DR_1455* | -1.669861229 | -2.130192143 | Uncharacterized protein |
| *DR_1610* | 1.104716541 | 1.162047146 | 3-isopropylmalate dehydratase large subunit 1 |
| *DR_1772* | -1.177473069 | -1.278671333 | Uncharacterized protein |
| *DR_1803* | -1.608202809 | -1.526228019 | Uncharacterized protein |
| *DR_1839* | -1.069929399 | -1.07154035 | Peptidyl-prolyl cis-trans isomerase |
| **DR_1840** | -1.006010815 | -1.387512842 | Uncharacterized protein |
| **DR_1844** | -1.435838931 | -1.287713524 | Uncharacterized protein |
| *DR_1886* | 1.00236572 | 1.15841242 | Uncharacterized protein |
| *DR_1914\** | 1.495789804 | -1.187629819 | Uncharacterized protein |
| **DR_2086** | -1.055246615 | -1.863947193 | Uncharacterized protein |
| *DR_2097* | -1.222613654 | -1.192798401 | Uncharacterized protein |
| *DR_2142* | 1.14465716 | 1.271880222 | Uncharacterized protein |
| *DR_2208* | 2.543721496 | 1.249885154 | Lactoylglutathione lyase, putative |
| *DR_2255* | 2.209610281 | 1.509724453 | Uncharacterized protein |
| *DR_2311* | 1.848131657 | 2.162303983 | Pseudouridine-5'-phosphate<br>glycosidase |
| *DR_2312* | 2.23497633 | 2.356794935 | Carbohydrate kinase, PfkB family |
| *DR_2362* | -1.165756078 | -1.02563511 | Transcriptional regulator |
| *DR_2519* | 2.37912365 | 1.286535846 | Transcriptional regulator, MerR<br>family |
| *DR_2559* | -1.091679205 | -1.606339412 | Uncharacterized protein |
| *DR_A0074* | 1.112580391 | 1.049621736 | Uncharacterized protein |
| *DR_A0075* | 2.982875996 | -1.364518319 | Transposase, putative |
| *DR_A0165* | 1.647644332 | 1.249869433 | Uncharacterized protein |
| *DR_A0218* | 1.44400857 | 1.056364322 | Uncharacterized protein |
| *DR_A0234* | -1.035264577 | -1.32234834 | Uncharacterized protein |
| *DR_A0263* | 1.85301791 | 1.260837031 | Branched-chain amino acid ABC<br>transporter, periplasmic amino<br>acid-binding protein |
| *DR_A0350* | -1.588574764 | -3.562978338 | Response regulator PilH, putative |
| *DR_A0352* | -1.426582502 | -4.426296882 | Methyl-accepting chemotaxis<br>protein |
| *DR_A0365* | 1.221175868 | 1.469528027 | Uncharacterized protein |
| *DR_B0016* | 2.093318651 | 1.210041621 | Hemin import ATP-binding<br>protein HmuV |
| *DR_B0038* | -1.225015351 | -2.813001105 | Uncharacterized protein |
| *DR_B0039* | -1.08507696 | -2.499499027 | Uncharacterized protein |
| *DR_B0054* | -1.133097457 | -1.213937901 | Uncharacterized protein |
| *DR_B0064* | -1.232475668 | -4.552778212 | Uncharacterized protein |
| *DR_B0122* | 2.417124882 | 1.093005808 | Iron ABC transporter, permease<br>protein |
| ***DR_C0001*** | -1.025302581 | -1.986341686 | Cytochrome P450-related protein |
| *DR_C0016* | -2.49574189 | -3.212004297 | Uncharacterized protein |
| ***DR_C0034*** | -1.329367995 | -1.640538179 | Coenzyme PQQ synthesis<br>protein, putative |
| *DR_t25\** | 1.031082599 | -1.177953562 | tRNA |

Moreover, although the large majority of differentially expressed genes were protein-coding genes, a few sRNAs and tRNAs were also impacted by the deletion of *khpA* or *khpB* (**Table S3**). Among the sRNA detected in our RNA-seq data (**Figure 5D**), *Dsr5*, *Dsr18*, *Dsr30* and *Dsr35*, *Dsr51* were significantly downregulated (|log2 fold change| ≥ 1, adjust *p* value ≤ 0.05) in the Δ*khpA* and/or Δ*khpB* strain compared to the WT, respectively. The deletion of KhpB also caused an increased level of *Dsr11*, consistent with our earlier observation that *Dsr11* is more stable in the absence of *khpB* (**Figure 3E**). Other sRNAs (such as *Dsr3*, *Dsr6*, *Dsr50*, *etc.*) showed differential expression in either of the KhpA or KhpB deletion strains, but these changes were less pronounced or less statistically significant (|log2 fold change| < 1, or adjust *p* value > 0.05). These data imply that KhpA and KhpB may affect the expression/stability of a wider range of sRNAs beyond those specifically examined in this study.

### KhpA and KhpB have no effect on cell division in *D. radiodurans*

Depletion of KhpA or KhpB has previously been shown to impair growth and alter cell division in *C. difficle*, *F. nucleatum*, *L. plantarum* and *S. pneumonie* (14, 15, 17, 18). However, we didn’t detect any significant changes in the abundance of genes known to be responsible for cell division of *D. radiodurans* (e.g. *DR_1369/divIVA*, *DR_0630/FtsA*, *DR_0631/FtsZ*, *DR_0379/BamA*, *DR_0400/FtsK*) between the WT and Δ*khpA* or Δ*khpB* strains. This observation implies that the KhpA and KhpB proteins may have different effects on cell size of *D. radiodurans*. To investigate this, we compared the growth of the WT and Δ*khpA*, Δ*khpB*, Δ*khpA*Δ*khpB* strains in TGY media and their cell size at mid-exponential phase. The depletion of *khpA*, *khpB*, or both from the *D. radiodurans* genome did not affect bacterial growth, although the mutant strains reached slightly lower final optical densities (1.13-1.21) compared to the WT (1.29) when entering stationary phase (**Figure S4C**). Moreover, the genomic deletion of *khpA* and *khpB* did not significantly alter the cell size of *D. radiodurans* under these growth conditions (**Figure S4D-S4E**). These results suggest the roles of KhpA and KhpB proteins in *D. radiodurans* differ from those in other bacteria, particularly in terms of bacterial growth and cell size.

### KhpA and KhpB affect expression of mRNAs targeted by sRNAs in D. radiodurans

Our data have suggested that *D. radiodurans* KhpA and KhpB can directly bind several sRNAs and can affect sRNA stability. However, the full scope as to how these proteins affect sRNA regulation (i.e. direct binding with their targets) is elusive across bacteria. To better understand the functional impact of KhpA and KhpB on sRNA biology in *D. radiodurans*, we analyzed the effects of their deletion on the expression of mRNAs previously shown to co-purify with *PprS* (29). We found that the expression of many RNAs associated with *PprS* (58 of 135) were altered by the deletion of KhpA and/or KhpB in *D. radioduran*s relative to the WT strain (**Table S4**). Importantly, the observed expression changes of RNAs affected by both proteins showed consistent trends (up- or down-regulated in both deletions), suggesting that KhpA and KhpB may cooperatively affect activity of *PprS* on these RNAs. Although we lack evidence that all of these RNAs are direct targets of *PprS*, their co-precipitation with the sRNA suggests that they could directly and/or indirectly physically associate with *PprS*. Based on the latter assumption, the overlap between the RNAs co-precipitated with *PprS* and those whose cellular abundance is impacted by KhpA or KhpB raise the possibility that KhpA or KhpB may impact sRNA biological activities directly or indirectly (i.e. via their interactions within protein networks). To further investigate whether KhpA or KhpB influence sRNA-target interaction, we performed a microscale thermophoresis (MST) assay. In this experiment, *pprM*, the characterized mRNA target of *PprS*, was labelled and incubated with *PprS* in the presence or absence of purified KhpA or KhpB proteins. We observed that the addition of either KhpA or KhpB protein remarkably enhanced the binding of *PprS* to *pprM* when compared to a control reaction without these proteins (**Figure 6A**). This data supports the hypothesis that KhpA and KhpB can influence sRNA-mRNA interaction, further suggesting that the presence of these proteins could play a role in modulating sRNA-mediated regulation of gene expression in *D. radiodurans*.

**Figure 6.**
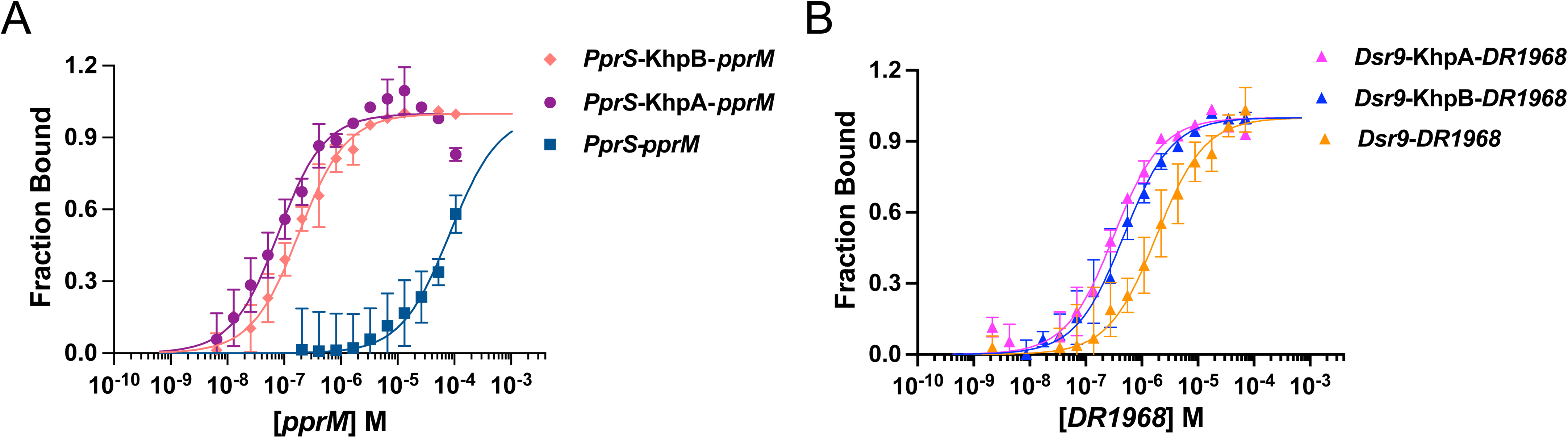

To test the generality of KhpA and KhpB effects on sRNA-mRNA regulation beyond *pprS*, we also examined the impact of these proteins on an additional sRNA, *Dsr9*. This sRNA was originally identified for its role in the biosynthesis of silver nanoparticles in *D. radiodurans* (28). However, we recently uncovered its contribution to the oxidation stress response. Upon knockdown of this sRNA led to reduced survival under oxidative stress conditions (**Figure S5A-S5B**). Importantly, we predicted *DR_1968*, a nitroreductase-encoding gene, as its primary target (**Figure S5C**). EMSA assays confirmed that *Dsr9* directly binds *DR_1968* mRNA within its coding region (**Figure S5D-S5E)**. qPCR analysis further suggested a negative regulation of *Dsr9* on *DR_1968* (**Figure S5F**). We then used this *Dsr9*-*DR_1968* pair to test the effect of KhpA and KhpB by MST analysis, which revealed that both proteins enhance the binding between *Dsr9* and *DR_1968* (**Figure 6B**). Altogether, these results demonstrate that the effects of KhpA and KhpB on sRNA-mRNA target interactions in *D. radiodurans* is not limited to the *PprS*-*pprM* pair.

## DISCUSSION

Despite the growing knowledge of biological roles of canonical bacterial sRNA binding proteins (Hfq, CsrA and ProQ), these proteins are not universally conserved across all bacteria (e.g., *D. radiodurans*). Recent studies strongly suggest that sRNA regulation and RBPs are key determinants of gene regulation in *D. radiodurans* (24–27, 33). However, although *D. radiodurans* encodes a plethora of regulatory RBPs, it lacks homologues of well-known Csr/Hfq/ProQ-like proteins, making it unclear as to which classes of proteins (if any) support sRNA function in this bacterium.

Echoing recent findings in several other gram-positive bacteria, such as *C. difficle and F. nucleatum* (17, 18), here we demonstrated that the KhpA and KhpB proteins in *D. radiodurans* associate with sRNAs that play essential roles in stress response (**Figure 2**). The sRNA binding ability of KhpA and KhpB in *D. radiodurans* is similar to that observed in *F. nucleatum* (in the micromolar range) (**Table 1**), indicating that these two RBPs act as weak sRNA binding proteins compared to Hfq/ProQ/CsrA, which usually exhibit low nanomolar binding affinity to their sRNA ligands (8, 36, 37). It is worth noting that weak RBPs with low RNA binding affinities are often implicated in RNA processing (i.e. through moonlighting activities) (40), suggesting that KhpA/B may have similar functions. Upon detailed investigation of the impact of KhpA and KhpB binding to four representative *D. radiodurans* sRNAs, we found that both proteins act as stability factors for at least some sRNAs. Similar results have been reported in *C. difficle and F. nucleatum* (17, 18). Interestingly, opposite effects were observed for these proteins in *D. radiodurans*: KhpA stabilized some of the tested sRNAs (*Dsr9*, *Dsr11*, and *Dsr20*), whereas the expression of KhpB destabilized *Dsr9* and *Dsr11* (**Figure 3**). Since KhpA and KhpB do not bind the same sRNA simultaneously as a complex (**Figure 4E-4H**), these two proteins presumably impact these sRNAs in different ways. We speculate that KhpB may recruit some ribonucleases and promote the degradation of some sRNAs, while the binding of KhpA with sRNAs protects them from the recognition of those ribonucleases. This idea is supported by the copurification of KhpB with Rsr (DR_1262) (**Table S2**), an RBP that recruits PNPase to promote RNA decay in *D. radiodurans* (41, 42). Notably, PNPase, which was previously shown to interact with sRNAs and modulate their turnover in both degradation and protection modes in *E. coli* (43), was also enriched in the 14-mer pulldown assay (**Figure 1C**). Although the specific ribonucleases regulating sRNA turnover in *D. radiodurans* remain undetermined, PNPase may mediate the stability of KhpA/KhpB-interacting sRNAs, either by itself or cooperatively with KhpA, KhpB and/or Rsr.

Through RNA-seq, we found that many mRNAs co-precipitated with *PprS* showed significant differential expression in the Δ*khpA* and Δ*khpB* strains (**Table S4**). This suggests that KhpA and KhpB could play important roles in sRNA activities, either directly or indirectly (i.e., through their network of interacting proteins). Interestingly, the most well-characterized *PprS* target (29), *pprM*, did not show any significantly differential expression in either Δ*khpA* or Δ*khpB*. However, we cannot rule out the possibility that KhpA and/or KhpB may contribute to the regulation of *PprS* on *pprM* under different (and specific) cell conditions, as the binding between *PprS* and *pprM* was stronger when incubating with KhpA or KhpB compared to the control (**Figure 6A**), and these proteins are essential for survival following radiation (30), a phenotype similar to the disruption of *PprS* or *pprM*. In contrast, the expression of DR_1968, the mRNA target of another sRNA *Dsr9*, was significantly upregulated when *khpA* was deleted (**Table S3**) and both KhpA and KhpB enhanced the *Dsr9*-*DR_1968* interaction (**Figure 6B**). These results imply that KhpA and KhpB may function as general sRNA-mRNA chaperones in *D. radiodurans*. Unfortunately, whether KhpA and KhpB directly bind more sRNAs in *D. radiodurans* cannot be concluded, since we failed to determine the direct RNA targetomes of KhpA and KhpB in this study, and many *D. radiodurans* sRNAs still remain uncharacterized. Nevertheless, given that several other sRNAs also showed differential expressions when KhpA or KhpB was deleted (**Figure 5D**), we speculate that KhpA and KhpB likely affect additional sRNAs and stress response in *D. radiodurans*.

Notably, it is intriguing to see that KhpA and KhpB interact with each other both *in vivo* and *in vitro*, but they do not bind same sRNAs as a complex in *D. radiodurans* (**Figure 4E-4H**). It is likely that they interact with these sRNAs individually when modulating their functions. This is consistent with our RNA decay experiments, which revealed opposing effects of KhpA and KhpB on sRNA stability (**Figure 3E**), further supporting their individual regulatory roles on sRNAs in *D. radiodurans*. The interaction between KhpA and KhpB with the sRNAs, as well as the interaction between the two proteins, may be mediated by the same residues on their KH domains, which was found to be crucial for the KhpA homodimerization and KhpA-KhpB heterodimerization in *S. pneumoniae* (38). This is supported by the observation of the RNA-independent interaction between KhpA and KhpB (**Figure 4C-4D**). However, in *F. nucleatum*, only KhpB forms complexes with sRNAs directly and this activity is enhanced by the addition of KhpA (17). Given that only a few sRNAs were tested in both studies, it is possible that only certain sRNAs bind to KhpA-KhpB protein complex, whereas others bind exclusively to one of the proteins.

Our work also reveals that KhpA and KhpB have no effect of on cell growth and size in *D. radiodurans* (**Figure S4C-S4E**), which is in stark contrast to previous findings in *C. difficle*, *F. nucleatum*, *L. plantarum* and *S. pneumonie* (14, 15, 17, 18). This discrepancy is probably related to the cellular localization of these two proteins. In *S. pneumoniae*, the KhpA-KhpB complex is recruited to the cell membrane by the transglycosylase MltG to regulate cell elongation in the division zone, a process mediated by the Jag-like domain on KhpB (38). Corroborating this, the KhpB protein of *F. nucleatum* was insoluble unless the Jag-like protein at the N-terminus was removed (17). However, this domain is missing in all of the *Deinococcus* KhpB homologs (**Figure S2D**) and we were able to purify this protein from the soluble fraction of *D. radiodurans* cell lysate, indicating a cytoplasmic location for KhpB in this bacterium. Additionally, it has been suggested that the localization of KhpB might be also linked to its upstream gene that encodes the YidC protein (18), which plays a central role in the insertion and/or folding of membrane proteins (44). However, both KhpA and KhpB exhibit different gene synteny in *D. radiodurans* compared to other bacteria (e.g., *S. pneumoniae*, *C. difficile* and *F. nucleatum*) (**Figure S2E-S2F**). These indicate a different cellular localization of KhpA or KhpB in *D. radiodurans* compared to other bacteria, which may explain why these proteins do not affect cell size in this bacterium.

In conclusion, this work advances our understanding of biological relevance of KhpA and KhpB in *D. radiodurans*. We anticipate future applications of RNA sequencing and structural analysis in combination with biochemical approaches will elucidate the full spectrum of RNAs that interact with KhpA and KhpB, how the two proteins recognize RNAs (e.g., through secondary RNA structures or motifs, what residues mediate their interaction with RNA targets), and the degree to which the RNA binding/regulatory activities of KhpA or KhpB rely on the interplay between the two proteins. We also hope that future studies will continue to elucidate how these mechanisms are conserved and vary across different classes of bacteria.

## MATERIAL AND METHODS

### Strains and cultural conditions

All strains used and constructed in this study are summarized in **Table S5**. The *D. radiodurans* R1 strain (ATCC 13939) is referred as wild-type (WT) throughout this study and was grown aerobically in TGY medium (1% tryptone, 0.1% glucose, 0.5% yeast extract) or on agar plates at 32°C. When needed, antibiotics were added at the following concentrations: chloramphenicol, 3.4 μg/mL; kanamycin, 16 μg/mL. *E. coli* strains were grown aerobically in Luria-Bertani (LB) medium (10 g/L tryptone, 10 g/L NaCl and 5 g/L yeast extract) or on agar plates at 37°C, supplemented with carbenicillin (50 μg/mL) or kanamycin (50 μg/mL).

### Plasmid and strain construction

Deletion mutants of *khpA* and *khpB* (Δ*khpA* and Δ*khpB*) were constructed via homologous recombination as previously described (27). Briefly, ∼1.2 kb of homology to the chromosomal sequence flanking the upstream and downstream regions of the *khpA* and *khpB* genes were amplified by PCR and assembled with a HindIII-digested pUC19mPheS plasmid, along with a kanamycin cassette using the NEBuilder HiFi DNA Assembly Master Mix (New England Biolabs). 1.5 µg of recombined pUC19mPheS plasmids were mixed with 955 μL of *D. radiodurans* WT cells at exponential phase (optical density at 600 nm (OD600) = 0.8), 45 μL of 1 M CaCl_2,_ and 500 μL of 30% glycerol, followed by 1 h incubation on ice and overnight incubation at 32°C. The next day, cells were plated on TGY plates with kanamycin and 5 mM 4-chloro-phenylalanine (Sigma-Aldrich). After 2-3 days, colonies grown on the plates were screened via colony PCR for insertion of the kanamycin resistance cassette and deletion of the target gene. The markless deletion strains were constructed by removal of the kanamycin cassettes. To achieve this, the pDeinoCre plasmid was transformed into the mutant strains, and the transformations were selected on both TGY plates with 5 mM 4-chloro-phenylalanine and TGY plates with 5 mM 4-chloro-phenylalanine and kanamycin. Those kanamycin-sensitive colonies were then screened by colony PCR to examine the loss of the kanamycin resistance cassette.

KhpB was tagged chromosomally with 3×FLAG or 6×His at the C-terminus by replacing the stop codon with a 3×FLAG or 6×His coding sequence using the same strategy as for gene deletion. However, the chromosomal tagging of KhpA cannot be achieved due to an unknown reason. Therefore, we amplified the coding sequence of KhpA (without stop codon) using a primer pair containing a 3×FLAG sequence at the 5’ end of the reverse primer so that 3×FLAG could be added to the C-terminus of KhpA. Tagged or untagged KhpA fragments were then ligated into the SacII/BamHI restriction sites on the pRADgro plasmid. The recombinant plasmids were transformed into the WT, Δ*khpA*, or the 6×His tagged KhpB strain, and the resulting colonies were selected on TGY plates containing chloramphenicol.

The complementary strains were constructed by cloning *khpA* or *khpB* coding sequence into the SacII/BamHI restriction sites on the pRADgro plasmid, after which the resulting constructs were transformed into the corresponding mutant strains. The construction of plasmids for protein purifications was achieved by cloning the coding regions of KhpA and KhpB proteins into the NdeI/BamHI restriction sites on the pET28a plasmid. The resultant plasmids were then transformed into the *E. coli* BL21 (DE3) strains, and the colonies were selected on LB agar containing kanamycin and verified by colony PCR.

All plasmids and primers are listed in **Table S5**.

### Protein extraction

The *D. radiodurans* WT, Δ*khpA,* and Δ*khpB* strains were grown in TGY until OD600 reached 0.8. Cells were collected from 10 mL of each culture and harvested by centrifugation at 4,000 rpm and 4°C for 10 min. The pellet was resuspended in 500 μl of 1×PBS supplemented with 1 mM PMSF (phenylmethylsulfonyl fluoride; Sigma-Aldrich) and lysed using a sonicator (XL-2000 Microson ultrasonic liquid processor; QSonica) for 10 cycles of 30 s bursts with 1 min of rest on ice between each burst. Following sonication, the samples were centrifuged at 12,000 rpm and 4°C for 10 min to get the soluble protein lysate in the supernatant. The protein concentration was determined using the Bradford assay.

### MS2-sRNA affinity purification

MS2-sRNA affinity pulldown was performed as described previously to determine the possible proteins interacting with *PprS in vivo* (11). The MS2 aptamer was fused to the 5’ end of the *PprS* sequence and cloned into the pRADgro plasmid, which was then transformed into WT *D. radiodurans*. A WT *D. radiodurans* strain expressing untagged *PprS* on the same vector was used as the negative control. Strains expressing these plasmids were cultured in TGY to mid-exponential phase (OD600 = 0.8) and cells were collected by centrifugation for at 4,000 rpm 10 min at 4°C,and then snap frozen. The cells were resuspended in 500 µL binding buffer [20 mM Tris-HCl (pH 8.0), 150 mM KCl, 1 mM of MgCl_2_, 1 mM DTT] supplemented with 1 mM PMSF and 0.4 U/mL SUPERase In RNase Inhibitor (ThermoFisher Scientific). The cells were mixed with 300 µL of 0.1 mm acid-washed glass beads (Sigma-Aldrich), chilled on ice for 3 min, and lysed through two rounds (100 s each) of bead-beating at 4°C. The lysates were cleared by centrifugation for 20 min at 13,000 rpm and 4°C. The supernatant was then incubated with 20 pmol purified MS2-MBP coat protein for 1 h at 4°C with rocking. The mixture was further incubated with amylose beads (New England Biolabs, pre-washed with 500 µL binding buffer) at 4°C for 2 h with rocking. The beads were pulled out by a magnet and washed with binding buffer five times. Bound proteins were eluted using 300 μL elution buffer (binding buffer with 15 mM maltose). After the addition of 1 mL Trizol reagent (ThermoFisher Scientific) and 300 µL of chloroform:isoamyl alcohol mix (v/v 24:1), the mixture was incubated at room temperature for 3 min and centrifuged at 13,000 rpm for 20 min at 4°C. 1.5 mL isopropanol was added to the phenol-chloroform layer, incubated for 10 min at room temperature, and centrifuged at 13,000 rpm for 30 min at 4 °C. The pellet was subsequently washed with 2 mL 0.3 M guanidine hydrochloride in 95% ethanol at room temperature three times, and 2 mL 100% ethanol twice. The protein pellets were air dried for 5 min and resuspended in 30 μL 1× protein loading buffer [0.5 M Tris-HCl, (pH 6.8), 25% (v/v) glycerol, 0.5% (w/v) SDS, 0.5% (w/v) Bromophenol blue, 0.05% (v/v) β-mercaptoethanol], which was boiled at 95 °C for 10 mins and prepared for LC-MS/MS analysis (described below). This assay was performed in two biological replicates.

### *In vitro* transcription and purification of RNAs

In *vitro* transcription was performed in 20 µL reactions using the Invitrogen MEGAscript T7 Transcription Kit (ThermoFisher Scientific) according to the manufacturer’s protocol. The RNA product was subjected to a denaturing urea PAGE with 6% polyacrylamide and 7 M urea. Gels were stained for 10 min by SYBR Gold (ThermoFisher Scientific), and the desired bands were cut out into small pieces and transferred into 2 mL tubes. 750 µL RNA elution buffer (0.1 M NaAc, 0.1% SDS, 10 mM EDTA) supplemented with 0.4 U/mL SUPERase In RNase Inhibitor was added to the tubes. After incubation overnight at 4°C with rocking, gel pieces were removed by centrifugation at 5,000 g and 4°C for 3 min. An equal volume of phenol:chloroform:isoamylalcohol (25:24:1, pH 4.5, Roth) was added to the supernatants, followed by centrifugation at 4°C and 13,000 rpm for 15 min. The aqueous phase was collected and subjected to ice-cold 30:1 ethanol: 3M NaOAc (pH 5.2) for overnight precipitation at - 20 °C. RNA was pelleted by centrifugation at 13,000 rpm for 30 min at 4°C and washed with ice-cold 70% ethanol, followed by centrifugation at 4°C and 13, 000 rpm for 10 min. The pellet was air-dried for ∼ 5 min and dissolved in 20 µL RNase-free water.

### 14-mer pulldown

The 14-mer pulldown was used as another method to identify *PprS*-associated proteins, based on a previously published method (32). *In vitro*-transcribed *PprS* containing a 14-mer tag was generated with a 39-nt 5’ overhang: GTTTTTTT<u>TAATACGACTCACTATA</u>**GGGAGACCTAGCCT**, where highlighted nucleotides represent the T7 promoter (underlined) and the 14-mer tag (bold). A scrambled RNA oligonucleotide with the same length as *PprS* (82 nt) containing the 14-mer tag served as the control. 4 µg of a 3’-biotinylated, 2’-O-methyl-modified RNA adapter complementary to the 14-mer tag on the *PprS* RNA (AGGCUAGGUCUCCC-biotin) was diluted in 500 µL binding buffer [50 mM Tris-HCl (pH 8.0), 150 mM KCl, 5 mM MgCl_2_, and 10% glycerol, 0.01% Tween-20] supplemented with 0.4 U/mL SUPERase In RNase Inhibitor and incubated with Dynabeads M-280 Streptavidin (ThermoFisher Scientific) for 1 h at 4°C. Next, the beads coupled with the adaptor RNA were applied to a magnet, washed twice, and resuspended in 1 mL binding buffer with SUPERase In RNase Inhibitor. 10 µg of 14-mer tagged *PprS* RNA was added to the beads, followed by a further incubation at 4°C for 2 h. *D. radiodurans* cells from a 500 mL culture grown to mid-exponential phase (OD600=0.8) were harvested and lysed in 800 µL binding buffer supplemented with 1 mM DTT and 1 mM PMSF by bead-beating. The lysate was centrifuged for 20 min at 4°C and 12,000 g to remove debris. Cell lysates were incubated with the coupled beads for 2 h at 4°C on a rotator in the presence of 0.4 U/mL SUPERase In RNase Inhibitor. The beads were washed once with ice-cold wash buffer 1 (binding buffer supplemented with 1 mM DTT, 1 mM PMSF, 300 mM KCl) and twice with ice-cold wash buffer 2 (binding buffer supplemented with 1 mM DTT, 1 mM PMSF). Finally, the beads were resuspended in 30 µL 1× protein loading buffer [0.5 M Tris-HCl, (pH 6.8), 25% (v/v) glycerol, 0.5% (w/v) SDS, 0.5% (w/v) Bromophenol blue, 0.05% (v/v) β-mercaptoethanol] and boiled at 95°C for 5 min. After 30 s of centrifugation at 500 rpm, the supernatant was transferred to a 1.5 mL tube for LC-MS/MS analysis. This assay was performed in two biological replicates.

### Protein purification

*E. coli* BL21 (DE3) carrying expression plasmids were grown in 2 L LB at 37°C to an OD600 nm of 0.6. IPTG was added to the cell culture a final concentration of 1 mM, followed by incubation at 30°C for 4 h with shaking. Cell pellets were then collected by centrifugation and resuspended in lysis buffer [50 mM Tris-HCl (pH 8.0), 300 mM NaCl, 10 mM imidazole, 1 mM PMSF]. The cells were then lysed through sonication for 20 cycles of 30 s bursts with 1 min of rest on ice between each burst. KhpA and KhpB proteins were subsequently purified from the soluble cell lysates using a Ni-NTA column (QIAGEN) as described previously (45). The proteins were finally concentrated with an Amicon centrifugal filter unit (3K cut-off, Millipore), and buffer exchanged to the protein storage buffer [100mM Tris-HCl (pH 8.0), 10% Glycerol, 1 mM DTT]. The quality of purified proteins was evaluated by SDS-PAGE and Coomassie staining, and the protein concentrations were determined by Bradford assay.

### Electrophoretic mobility shift assays

Electrophoretic mobility shift assays (EMSA) were performed to determine the binding affinity of KhpA and KhpB proteins to *D. radiodurans* sRNA *in vitro*. Selected sRNA candidates were *in vitro* transcribed, and gel purified as described above. 100 pmol of each RNA was then dephosphorylated by incubating with 25 U of calf intestinal alkaline phosphatase (New England Biolabs) at 37°C for 1 h. The dephosphorylated RNA was extracted and precipitated following the protocol outlined above. To prepare radiolabeled RNA, 50 pmol of each RNA were 5’ end-labeled using 50 µCi ^32^P-γATP (PerkinElmer) and T4 polynucleotide kinase (New England Biolabs) at 37°C for 1 h. The labeled RNA was then subjected to another round of gel extraction and eluted in 15 µL RNase-free water. For EMSA, 0.1 pmol of radiolabeled sRNAs were incubated with varying concentrations of purified KhpA and KhpB proteins (0∼10,000 nM) in 10 µL reactions containing 1× EMSA binding buffer [20 mM Tris-HCl (pH 8.0), 1 mM MgCl_2_, 20 mM KCl, 10 mM Na_2_HPO_4_-NaH_2_PO_4_ (pH 8.0), 10% glycerol, 0.4 U/mL SUPERase RNase Inhibitor]. To test interaction between *Dsr9* and *DR_1968*, DNA templates of *Dsr9* or *DR_1968* mRNA fragment (-100 bp to + 200 bp from the start codon) with or without designed mutations was synthesized (from Integrated DNA Technologies, Inc.). *In vitro* transcription and purification were carried out as described above. 1 pmol of radiolabeled *Dsr9/Dsr9\** was incubated with varying concentrations of *DR_1968*/*DR_1968\** (0∼18 nM) in 10 µL reactions containing 1× EMSA binding buffer. Reactions were incubated at 37°C for 1 h and resolved in 8% polyacrylamide native gels, followed by visualization on a Typhoon FLA 700 imager (GE Health Life Science). When needed, the band intensities were calculated using Image J, and the equilibrium dissociation constant (K_d_) was determined as described before (45). Each EMSA assay was conducted two times.

### Size exclusion chromatography

Purified KhpA or KhpB protein was mixed at a 1:1 molar ratio (15 nmol each) in a total volume of 300 µL in 1x EMSA binding buffer and incubated at 4°C for 1 h. The mixture was then loaded onto a Superdex 75 gel filtration column (Cytiva) that was pre-equilibrated in 1x EMSA binding buffer. The elution fractions were collected at a flow rate of 0.5 mL/min and the UV absorbance at 280nm was monitored throughout the run. For controls, individual proteins were loaded separately on the column under the same condition.

### Protein co-immunopurification

Protein co-immunopurification experiments were carried out as described previously (33). Strains expressing 3×FLAG tagged KhpA, 3×FLAG tagged KhpB, or 6×His tagged KhpB (**Table S5**) were grown in TGY medium to an OD600 of 0.8. 50 OD of each cell culture was harvested by centrifugation for 10 min at 4,000 rpm and 4°C, snap frozen, and resuspended in 800 µL lysis buffer 1 [20 mM Tris-HCl (pH 8.0), 1 mM MgCl_2_, 150 mM KCl, 1 mM DTT]. Subsequently, each sample was mixed with 1 U DNase I (Fermentas), 1 mM PMSF, 0.4 U/mL SUPERase In RNase Inhibitor (ThermoFisher Scientific) and 800 µl of 0.1 mm acid-washed glass beads (Sigma-Aldrich), and lysed by bead-beating at 4°C. For coIP using 3xFLAG tagged strains, 900 µL of clarified cell lysates were incubated with 25 µL of mouse-anti-FLAG antibody (Sigma-Aldrich) at 4°C for 1 h (rocking). Each sample was incubated with 75 µL protein A sepharose (Sigma-Aldrich), which has been pre-washed with 1 ml lysis buffer 3 times, for another hour at 4°C with rocking. Beads were washed 5 times with 500 µL lysis buffer and then resuspended in 500 µL of the same buffer. To elute coimmunoprecipitated proteins, 500 µL of Phenol:Chloroform:Isoamylalcohol (25:24:1, pH 4.5, Roth) was added, and the mixture was incubated at room temperature for 3 min. After centrifugation at 13,000 rpm and 4°C for 20 min, the organic phase was combined with 1.4 mL ice-cold acetone. After overnight precipitation at - 20°C, the tubes containing the beads and acetone were centrifuged at 12,000 rpm and 4°C for 60 min. The supernatant was carefully removed and the beads containing the coimmunoprecipitated proteins were washed twice with 1 mL of ice-cold acetone. The beads were then air-dried for 10 mins and resuspended in 50 μL of 1× protein loading buffer [0.5 M Tris-HCl, (pH 6.8), 25% (v/v) glycerol, 0.5% (w/v) SDS, 0.5% (w/v) Bromophenol blue, 0.05% (v/v) β-mercaptoethanol], boiled for 10 min at 95°C, and centrifuged for 5 min at 12,000 rpm and 4°C to collect the co-purified proteins. For coIP using 6×His tagged strains, 50 OD of cell culture was resuspended and lysed in lysis buffer 2 [50 mM Tris-HCl (pH 8.0), 1 mM MgCl_2_, 150 mM KCl, 1 mM DTT, 20 mM imidazole] and lysed as described above. 900 µL of clarified cell lysates were incubated with 25 µL of Dynabeads His-Tag Isolation and Pulldown magnetic beads (ThermoFisher Scientific) for 1 h at 4°C with gentle rocking. The beads were washed five times with lysis buffer 2, and co-immunopreciated proteins were eluted from the beads using the elution buffer [50 mM Tris-HCl (pH 8.0), 1 mM MgCl_2_, 150 mM KCl, 1 mM DTT, 200 mM imidazole]. The proteins from three independent coIP replicates were subjected to LC-MS/MS or Western blotting analysis. To examine if the protein-protein interactions are RNA-dependent, RNase Inhibitor was omitted in the coIP experiments, and a mixture of RNase A (2 μg) and RNase T1 (5 U; Thermo Fisher Scientific) was added to the cell lysates. The mixture was incubated for 15 min at 22°C before proceeding with antibodies or magnetic beads for further coIP steps.

### LC-MS/MS

The proteins obtained from pulldown/co-purification experiments were run ∼1 mm into a 12% SDS-PAGE gel. Gels were stained with Coomassie Blue, and the protein bands were cut off. The gel slices were then reduced with 10 mM DTT for 30 min at room temperature and alkylated using 50 mM iodoacetamide for 30 min at room temperature in the dark. The protein-containing gel pieces were digested with 10 ng/μl trypsin (ThermoFisher Scientific) overnight at 37°C. Proteins were extracted from the gel using 5% formic acid and 1:2 (v/v) 5% formic acid:acetonitrile. The obtained protein samples were sent to the University of Texas ICMB proteomics facility for LC-MS/MS analysis. The resulting protein spectral counts were searched against the *D. radiodurans* R1 (ATCC 13939) Uniprot database, and analyzed by Scaffold v4.4.1 (Proteome Software). A database of common contaminants was also included in the search. Protein identification was controlled using a false discovery rate (FDR) of 1 % at both the protein and peptide levels. Proteins identified with fewer than two unique peptides were dismissed. The difference between two strains/fractions was calculated using the Student’s t-test, with *p* values adjusted by the Benjamini-Hochberg method (46). Proteins with fold change of ≥1.5 or ≤0.67 and an adjusted *p* value of ≤ 0.05 were considered as significant differential expressed.

### Western Blotting

Samples from the coIP experiments were separated by 12% SDS-PAGE and transferred to nitrocellulose membranes (Bio-Rad) for 20 min at 4°C and 25 V. The membranes were blocked with 5% milk powder (in 1x TBS) for 1 h at room temperature. After washing three times with 1×TBST (1×TBS supplemented with 0.1% Tween 20) on a shaker (5 min each), the membranes were incubated overnight at 4°C with anti-FLAG (1:2,000) or anti-His antibodies (1:2,500) (Sigma Aldrich) in 3% milk powder (in 1×TBS). The membranes were washed again three times with 1×TBST. Next, 1 mL of ECL substrate (GE Healthcare) was added to the membrane and chemiluminescence was detected with a CCD camera (ImageQuant, GE Healthcare). Western blotting analysis of co-immunopurification fractions was performed in duplicate.

### RNA sequencing and data analysis

*D. radiodurans* WT, Δ*khpA,* and Δ*khpB* strains was inoculated in TGY medium to a final OD600 of 0.8. 10 mL of cells were harvested by centrifugation for 10 min at 4,000 g and 4°C. The cells were then mixed with 1 mL TRIzol reagent and lysed through two rounds of bead-beating on ice (100 s each). 400 µL chloroform was added to the lysate and mixed. After centrifugation for 15 min at 4°C and 13, 000 rpm, RNA was precipitated with ice-cold 30:1 ethanol: 3M NaOAc (pH 5.2), washed with ice-cold 70% ethanol twice, and dissolved in 35 µL RNase-free water. The RNA samples were then digested with DNase I (New England BioLabs) for 30 min at 37°C and purified using the RNA Clean & Concentrator Kit (Zymo Research). The RNA samples were quantified and evaluated using Bioanalyzer and Qubit 4 Fluorometer (ThermoFisher Scientific) before sequencing. Ribosomal RNA was depleted using the NEBNext rRNA Depletion Kit (Bacteria) (New England BioLabs). NEBNext Multiplex RNA Library Prep Set for Illumina (New England Biolabs) was used for library construction and sequencing was performed by the Genomic Sequencing and Analysis Facility at the University of Texas at Austin using the Illumina NovaSeq S1 single-end platform.

Approximately 10 million reads were mapped to the genome for each sample. The adapter sequences were removed using CutAdapt (47) and the reads shorter than 22 nt and/or with low-quality ends (phred quality < 30) were discarded. The remaining reads were mapped to the *D. radiodurans* R1 genomes (GenBank accession numbers NZ_CP038663.1. NZ_CP038664.1 NZ_CP038665.1 and NZ_CP038666.1) using BWA (48). Aligned reads were assigned to gene regions using HTseq. Differential expression of genes was normalized and calculated by the DEseq2 algorithm (49). Transcripts with log2 fold change of ≥1 or ≤-1 and an adjusted *p*-value of ≤ 0.05 were considered as significant differentially expressed. Each strain was analyzed in three biological replicates. Gene ontology (GO) analysis was performed through the David webtool (50).

### Northern Blotting

*PprS* expression was monitored by Northern blotting as described before (29). 5 μg of total RNA from each sample were loaded per lane and separated on 8% (vol/vol) polyacrylamide (PAA)-7 M urea gels in 1×TBE buffer at 15 W for 8 h. RNA was transferred onto Hybond X+ membranes (GE Life Sciences) by electroblotting at 15 V and 4°C for 16 h. The membranes with transferred RNA were then treated with UV light for crosslinking and hybridized overnight at 42 °C with *PprS*-specific radiolabeled ssDNA oligonucleotides (**Table S5**) at a final concentration of 100 nM in ULTRAhyb hybridization buffer (ThermoFisher Scientific). Following hybridization, the membranes were washed once with 5×SSC and 0.1% SDS at 30°C, then washed twice with 1×SSC and 0.1% SDS at 42°C. After exposure to a phosphorimaging screen for at least 6 h, the signals were read on a Typhoon FLA 700 phosphorimager (GE Healthcare) and quantified using the CLIQS software (TotalLab). Signal intensity for *PprS* was normalized to that of 5S rRNA which served as an internal loading control. Each experiment was performed in two biological replicates.

### Quantitative PCR (qPCR)

Total RNA samples were extracted and treated with DNase I as stated above. qPCR was performed using the Luna Universal One-Step RT-qPCR Kit (New England BioLabs) according to the manufacturer’s protocol on a ViiA7 instrument (Applied Biosciences). In each reaction, 25 ng of RNA was used as the template. The thermal cycling conditions included cDNA synthesis at 55°C for 10 min, initial denaturation at 95°C for 1 min, followed by 40 cycles of 95°C for 10 s and 60°C for 30 s. Relative fold change was calculated with the 2^−ΔΔCT^ method (51). Each qPCR experiment was performed in triplicate, and data were normalized to 16S rRNA (*DR_r01*). All primers used for qPCR are listed in **Table S5**.

### RNA decay assay

*D. radiodurans* WT, Δ*khpA*, and Δ*khpB* strains were grown to mid-exponential phase (OD600=0.8) in TGY media without antibiotics. Following the addition of rifampicin (VWR) to a final concentration of 250 μg/mL to abrogate transcription, samples were collected at indicated time points and snap-frozen in liquid nitrogen. RNA was extracted from those samples as described above and analyzed via Northern blotting or qPCR analysis. Calculation of half-lives was performed with GraphPad Prism version 9.0 by applying one phase decay equation. Each experiment was performed in three biological replicates.

### Survival assay

To test the impact of *Dsr9* on survivability of *D. radiodurans* under oxidative stress, biological triplicates of WT and *Dsr9* KD strains were grown in TGY media at 32°C to an OD600 of 0.8. 100 µL of each cell culture was added to sterile flat-bottom 96 well plates. Filter-sterilized H_2_O_2_ in 1×PBS was added to each well containing cell culture to reach final H_2_O_2_ concentrations of 0, 100 and 200mM. The cells were further incubated for 30 min at 4°C in the dark. Cell clutures were then immediately serially diluted in sterile 1×PBS (10^-0^ to 10^-4^), and 8 µL of each dilution were plated on TGY agar plates. Plates were grown at 32°C for 2 days and then imaged.

The alamarBlue assay was further carried out to compare the survival of WT and *Dsr9* KD under H_2_O_2_ stress. For this experiment, six biological triplicates of each strain were grown as described above and plated in 96 well plates for H_2_O_2_ exposures. Following the 30 min H_2_O_2_ exposure at 4°C in the dark, the plates were spun down at 4,000 rpm for 10 min at 4°C. The supernatant was removed, and cells were resuspended in 100 µL 10% alamarBlue (ThermoFisher Scientific) in TGY media. The cells were then incubated at 32°C in the dark for 1 h. Fluorescence (Excitation: 560nm; Emission: 590nm) was then measured using a plate reader (BioTek Cytation 3). The average fluorescence of wells containing only TGY media (background) was then subtracted from the experimental wells.

### Microscale Thermophoresis Assay

Binding assessment of *PprS/Dsr9* to *pprM/DR_1968* in the presence or absence of KhpA or KhpB were assessed through microscale thermophoresis (MST) using the Nanotemper Monolith instrument. sRNAs (*PprS* and *Dsr9)* and mRNA targets (*pprM* and *DR_1968)* were *in vitro* transcribed as depicted earlier in this manuscript. *PprM* or *DR_1968* was fluorescently labeled with Cy5 using the Label It Nucleic Acid Cy5 labeling kit (Mirus Bio) following the manufacturer’s instruction. Labeled RNA was purified using G50 microspin purification columns and total recovery was assumed as specified in the protocol. Monolith software was utilized to establish the dilution protocol for two-molecule binding and ternary complex binding. Briefly, fluorescently labeled *pprM* or *DR_1968* was diluted with binding buffer [20 mM Tris-HCl (pH 8.0), 1 mM MgCl_2_, 20 mM KCl, 10 mM Na_2_HPO4-NaH_2_PO_4_ pH 8.0, 10% glycerol, 0.05% Tween 20 and 1 mM DTT]. A serial dilution of *pprM* or *DR_1968* was prepared, ranging from 105.5 to 3.21x10^-3^ μM final concentration, and mixed with either *PprS or DR_1968* alone (20 nM) or with *PprS or Dsr9* and KhpA (2 μM) or KhpB (2 μM) in binding buffer. The samples were incubated at room temperature for 30 min and then loaded into nanotemper premium capillaries. Fluorescent measurements were made at 50% excitation and medium laser power. Data analysis was performed using the software’s K_d_ model, with data normalized from 670 nm/650 nm ratios to fraction bound for comparison purposes. These assays were conducted in three biological replicates.

### Growth curve measurement

Growth curves of *D. radiodurans* strains were measured using a Plate Reader (BioTek Cytation 3). Overnight cultures were diluted in biological triplicates to an initial OD600 of 0.1 in TGY media. 200 μL cell cultures of each strain were added to 96-well plates. The turbidity at 600 nm was measured every 15 min for 24 h with shaking at 200 rpm at 32°C.

### Microscope analysis

Cell cultures were grown to mid-exponential phase (OD600=0.8) in TGY media. 1 mL of bacterial cells suspended in 1 mL sterile 1×PBS were incubated with Nile Red (ThermoFischer Scientific) at a final concentration of 30 µM for 15 min at 32°C with agitation. The cells were collected by centrifugation for 1 min at 3,000 rpm, resuspended again in 200 µL sterile 1×PBS, and loaded onto a microscope slide topped with a coverslip. Images were taken with an Olympus FV1000 motorized inverted IX81 microscope using a 488nm laser. Cell volumes were measured using an in-house Python script. More than 400 cells from three biological replicates were measured. Statistical significance was determined by one-way ANOVA analysis (using GraphPad Prism).

### Statistics analysis

Data are expressed as the means ± SEMs. The Student’s t-test, one-way or two-way ANOVA analysis was used to evaluate the significance. *P* value cutoffs for significance were ≤ 0.05 (*), ≤ 0.01 (**), ≤ 0.001 (***), and ≤ 0.0001 (****).

## Data Availability

All RNA-sequencing data are available in the SRA database and can be publicly accessed via BioProject number PRJNA948458. All LC-MS/MS raw data have been deposited to the ProteomeXchange Consortium via the PRIDE partner repository with the dataset identifier PXD052534 (MS2-sRNA affinity pulldown), PXD052535 (14 mer pulldown), and PXD052557 (protein coIP). The python script for microscopy analysis has been made publicly available in Zenodo (https://doi.org/10.5281/zenodo.7787206).

## ACKNOWLEDGEMENTS

This work was supported by the National Institute of Health [R01GM135495 to L.M.C.; R01AI121500 to V.G.], Welch Foundation [F-1756 to L.M.C.], the Defense Threat Reduction Agency Young Investigator Program [HDTRA1-17-1-0025 to L.M.C.], the Air Force Office of Scientific Research Young Investigator program [FA9550-13-1-0160 and FA9550-20-1-0131 to L.M.C.] and NSF Civil, Mechanical and Manufacturing Innovation (CMMI) award [2150878, to V.G.]. The funding agencies had no role in the design of the study and collection, analysis, and interpretation of data or in writing the manuscript.

R.H. designed all experimental work, performed the majority of the experimental work, wrote and revised the manuscript. R.H. and J.F. constructed all the strains. R.H., B.Q.D., and J.F. performed the qPCR and Northern blotting experiments. B.Q.D. performed the MST experiments. B.N., A. Cordova., and V.G. designed and performed the microscope analysis. A. Cordova., S.E. and A. Chen performed the experiments for Dsr9 characterization. R.H. and P.S. analyzed the RNA sequencing data. R.H. collected and analyzed all the data. R.H. and L.M.C. supervised this work and revised the manuscript. All authors read and approved the final manuscript.

We thank the ICMB proteomics facility at the University of Texas at Austin for performing the LC-MS/MS analysis, and the Genomic Sequencing and Analysis Facility at the University of Texas at Austin for performing the RNA sequencing. We would also like to acknowledge all the Contreras lab members who provided helpful conversations and insights.

We declare that there are no conflicts of interest.

---

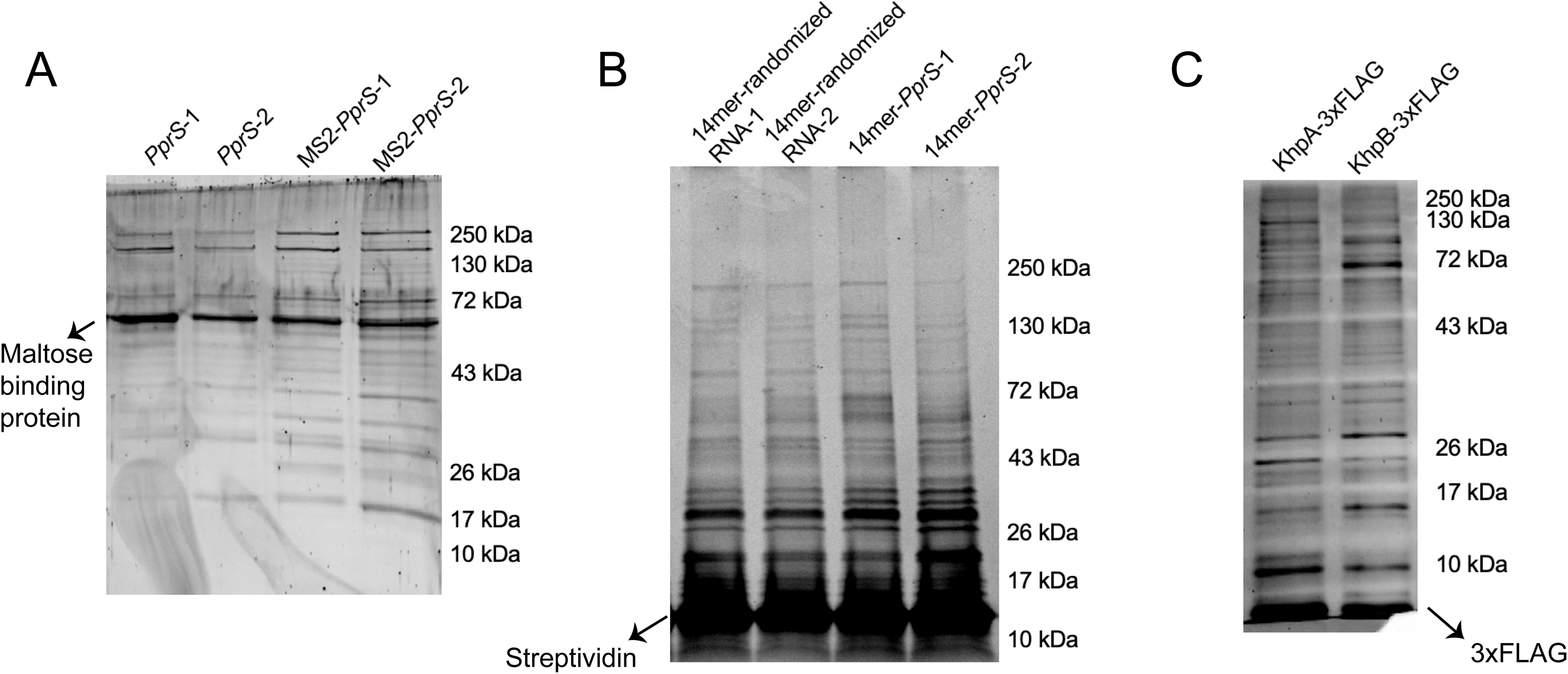

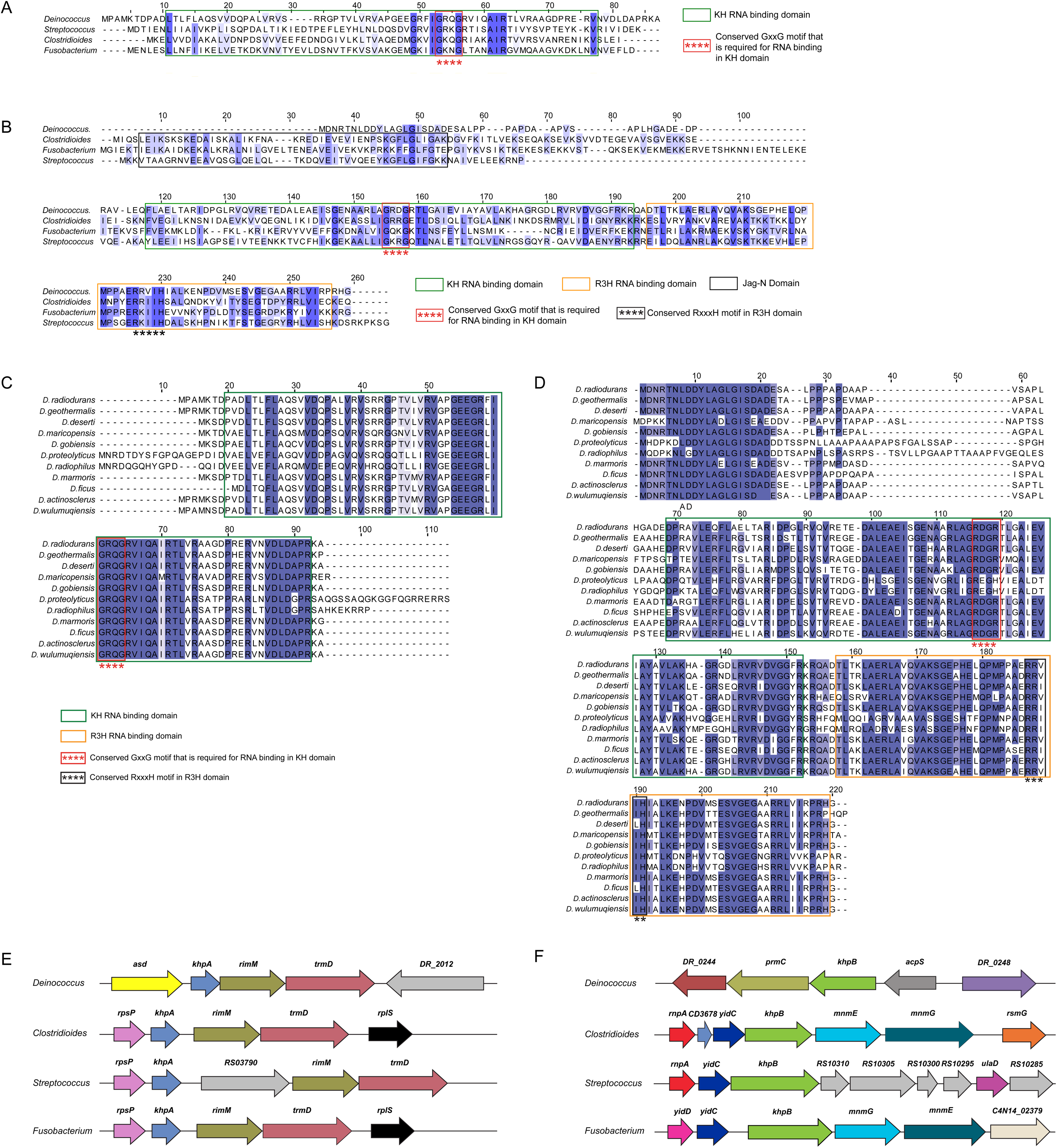

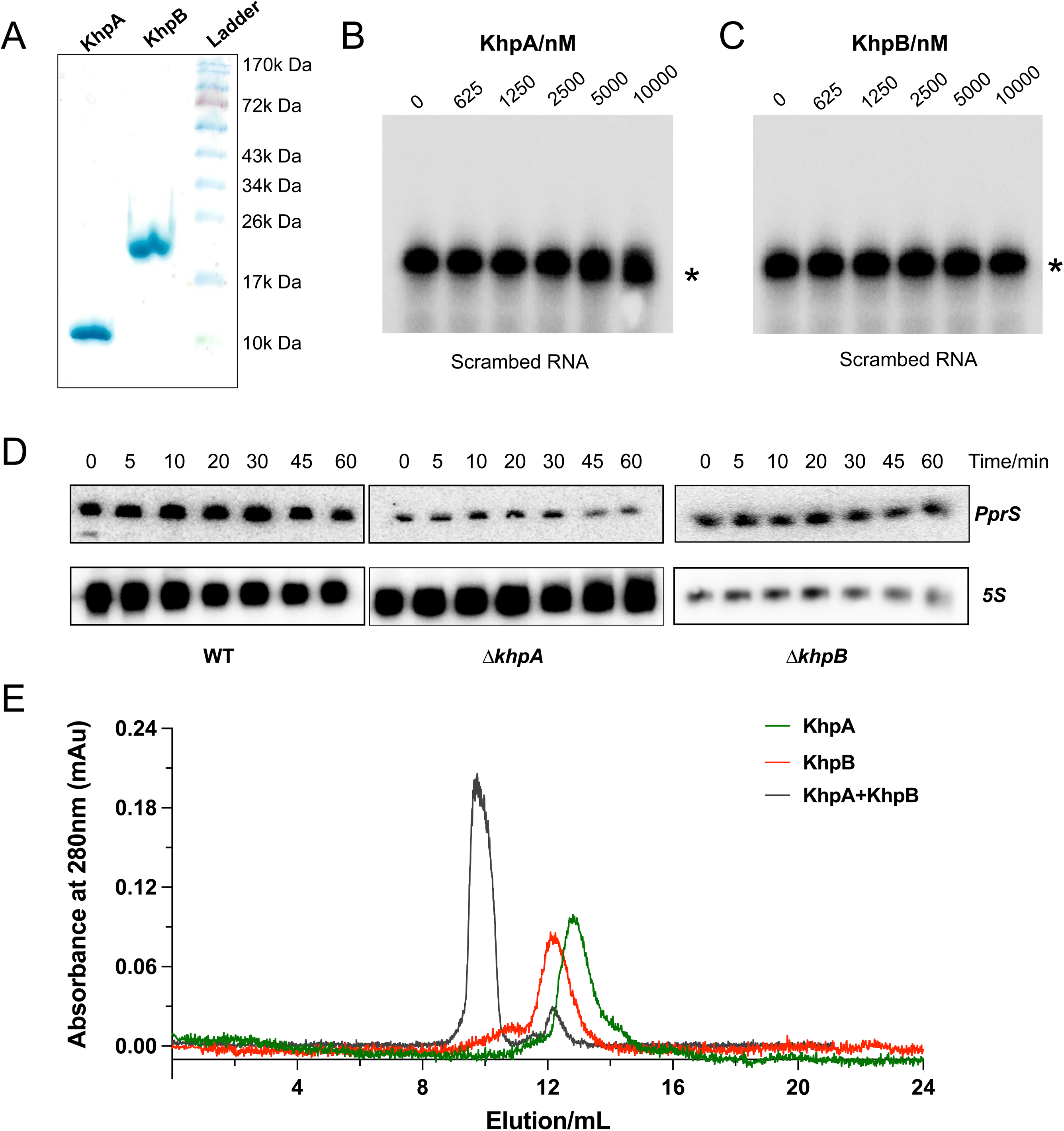

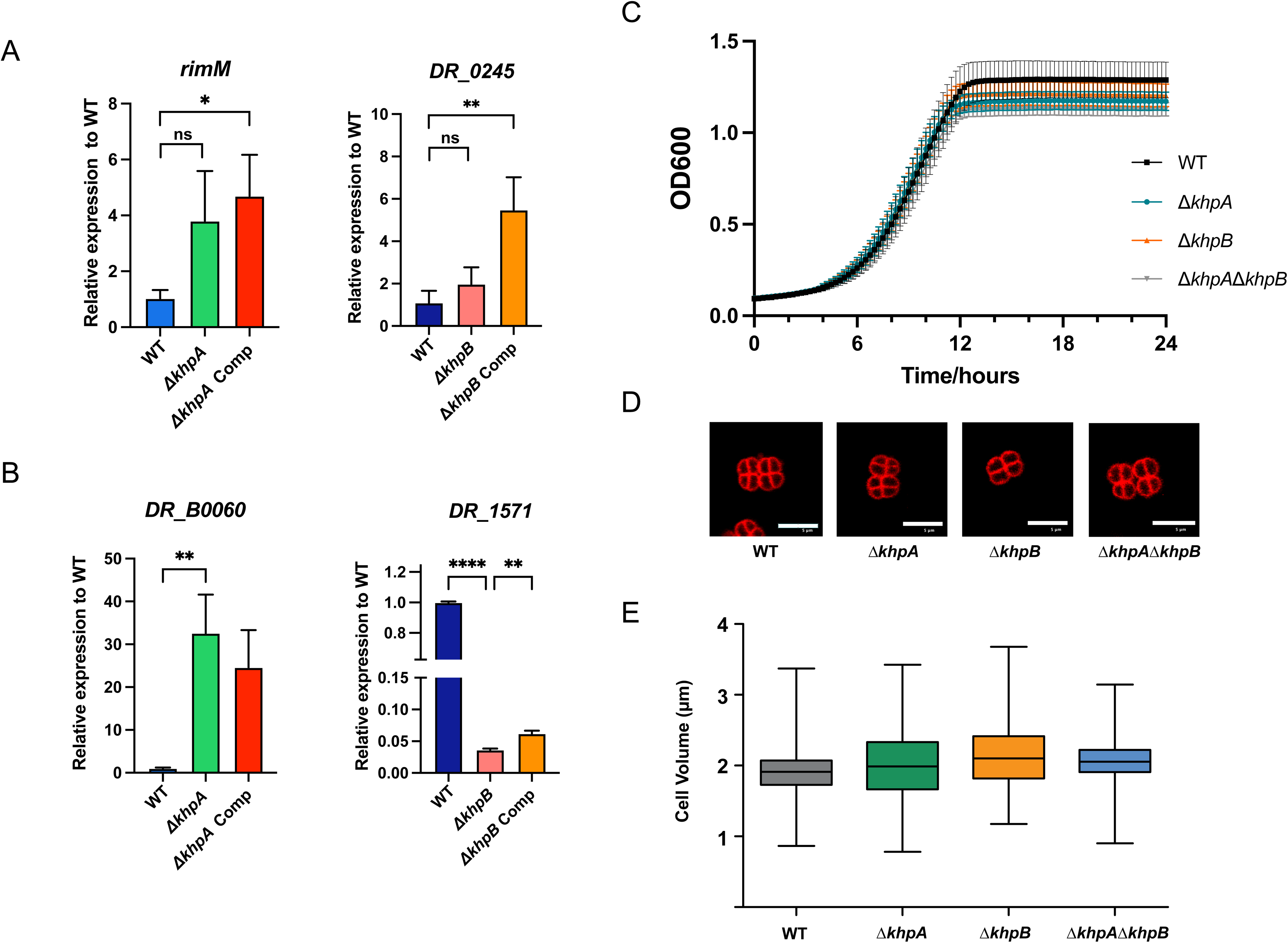

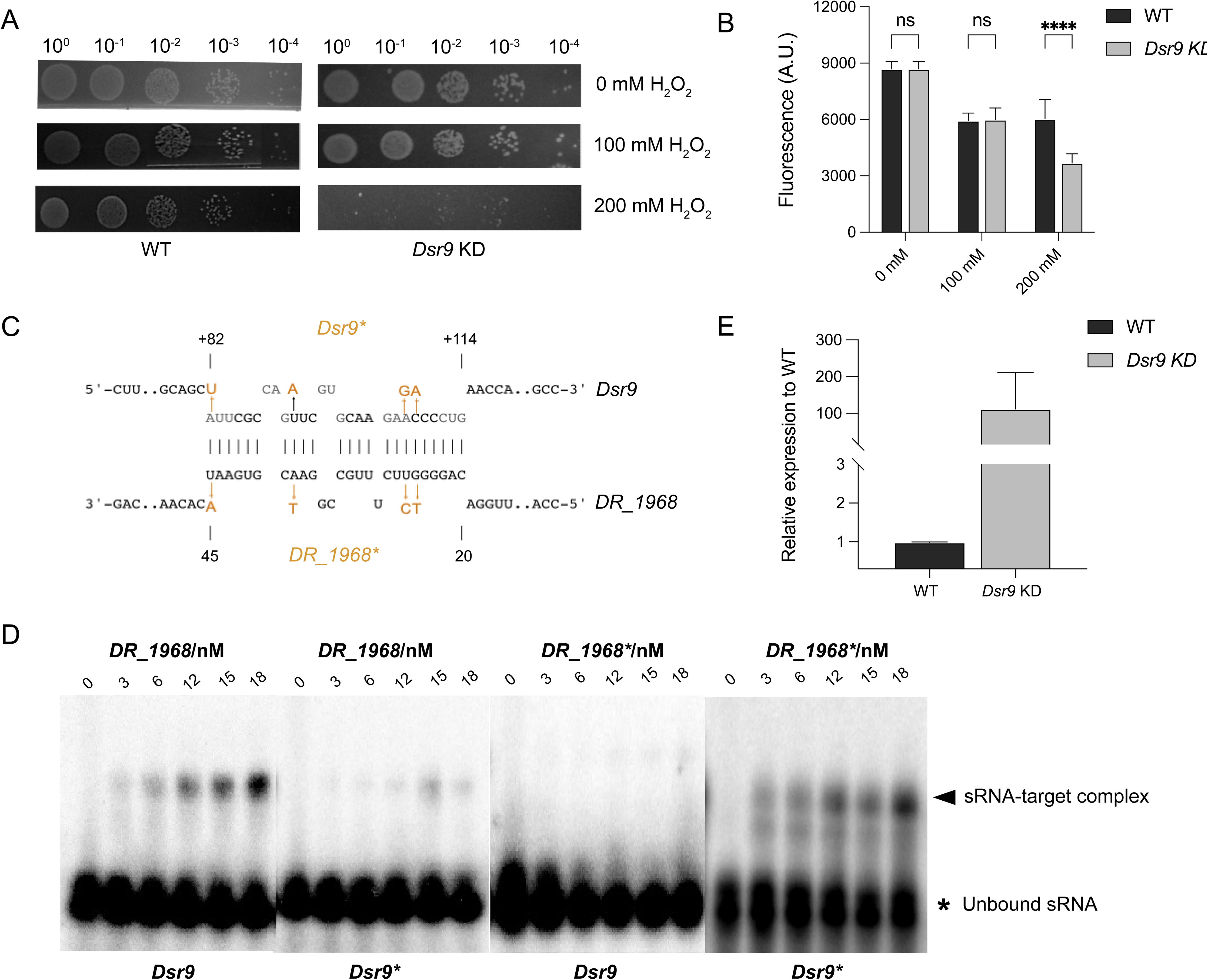

